# Single cell multi-omics profiling reveals a hierarchical epigenetic landscape during mammalian germ layer specification

**DOI:** 10.1101/519207

**Authors:** Ricard Argelaguet, Hisham Mohammed, Stephen J Clark, L Carine Stapel, Christel Krueger, Chantriolnt-Andreas Kapourani, Yunlong Xiang, Courtney Hanna, Sebastien Smallwood, Ximena Ibarra-Soria, Florian Buettner, Guido Sanguinetti, Felix Krueger, Wei Xie, Peter Rugg-Gunn, Gavin Kelsey, Wendy Dean, Jennifer Nichols, Oliver Stegle, John C Marioni, Wolf Reik

## Abstract

Formation of the three primary germ layers during gastrulation is an essential step in the establishment of the vertebrate body plan. Recent studies employing single cell RNA-sequencing have identified major transcriptional changes associated with germ layer specification. Global epigenetic reprogramming accompanies these changes, but the role of the epigenome in regulating early cell fate choice remains unresolved, and the coordination between different epigenetic layers is unclear. Here we describe the first single cell triple-omics map of chromatin accessibility, DNA methylation and RNA expression during the exit from pluripotency and the onset of gastrulation in mouse embryos. We find dynamic dependencies between the different molecular layers, with evidence for distinct modes of epigenetic regulation. The initial exit from pluripotency coincides with the establishment of a global repressive epigenetic landscape, followed by the emergence of local lineage-specific epigenetic patterns during gastrulation. Notably, cells committed to mesoderm and endoderm undergo widespread coordinated epigenetic rearrangements, driven by loss of methylation in enhancer marks and a concomitant increase of chromatin accessibility. In striking contrast, the epigenetic landscape of ectodermal cells is already established in the early epiblast. Hence, regulatory elements associated with each germ layer are either epigenetically primed or epigenetically remodelled prior to overt cell fate decisions during gastrulation, providing the molecular logic for a hierarchical emergence of the primary germ layers.

**Highlights:**

- First map of mouse gastrulation using comprehensive single cell triple-omic analysis.
- Exit from pluripotency is associated with a global repressive epigenetic landscape, driven by a sharp gain of DNA methylation and a gradual decrease of chromatin accessibility.
- DNA methylation and chromatin accessibility changes in enhancers, but not in promoters, are associated with germ layer formation.
- Mesoderm and endoderm enhancers become open and demethylated upon lineage commitment.
- Ectoderm enhancers are primed in the early epiblast and protected from the global repressive dynamics, supporting a default model of ectoderm commitment *in vivo*.

---

In mammals, specification of the basic body plan occurs during gastrulation, when a single-layered blastula is reorganised to give rise to the three primary germ layers: the ectoderm, mesoderm and endoderm^1,2^. Recent advances in single-cell RNA sequencing have facilitated unbiased and comprehensive characterisation of such developmental trajectories in a variety of different systems^3–5^. However, while gene expression dynamics of mouse development have been characterized in detail^6–12^, the role of the epigenome in cell fate decisions in early development remains poorly understood^13^.

The period of mouse development prior to gastrulation is characterised by extensive remodelling of the epigenome^14–25^. Moreover, recent studies have identified cell type specific chromatin marks, accessibility and DNA methylation profiles at regulatory elements in several species^26–31^. Together, this suggests that correct establishment of chromatin accessibility and DNA methylation may be important for cell fate specification during development. Indeed, mutants that fail to correctly remodel their epigenetic landscape display differentiation defects at or following gastrulation^32,33^. However, since cell fate decisions are made at the level of single cells, the ability to understand whether distinct epigenetic environments precede or follow cell fate choice during early development has been challenging. Moreover, in isolation, characterising the epigenome of individual cells provides only part of the picture – without knowledge of the transcriptional readout, associating epigenetic changes with specific cell fate choices is difficult^13^.

Recent technological advances have enabled the profiling of multiple molecular layers in single cells^34–43^, providing novel opportunities to study the relationship between the transcriptome and epigenome during cell fate decisions. Here we apply scNMT-seq (single-cell Nucleosome, Methylome and Transcriptome sequencing) to comprehensively analyse the collective dynamics of RNA expression, DNA methylation and chromatin accessibility in peri-implantation and early postimplantation mouse embryos during exit from pluripotency and primary germ layer specification.

## Mapping RNA expression, DNA methylation and chromatin accessibility changes during mouse gastrula development

We applied scNMT-seq^34^ to jointly profile chromatin accessibility, DNA methylation and gene expression from 758 single cells isolated from mouse embryos at four developmental stages (Embryonic Day (E) 4.5, E5.5, E6.5 and E7.5) spanning implantation and gastrulation **(Figure 1a-d)**. Following quality control and exclusion of trophoblast cells, 687 cells remained for downstream analysis (**Figure S1**).

As previously described^7–9^ we observe an increase in transcriptional diversity as embryos transition from a relatively homogeneous blastula to a heterogeneous gastrula with the ingression of cells through the primitive streak and the emergence of the three germ layers (**Figure 1a,b**). Gene expression profiles separate cells from the extraembryonic endoderm (E4.5 to E6.5), epiblast (E4.5 to E6.5), primitive streak (E6.5) and the three germ layers (E7.5) (**Figure S2-S5)**, consistent with published bulk-RNA-seq data from the same stages^16^ (**Figure S6**). Similarly, dimensionality reduction of DNA methylation and chromatin accessibility data reveals separation of cells by stage (**Figure 1c,d**) and embryonic versus extraembryonic origin (**Figure S7,S8**). These observations are consistent with previously published bulk DNA methylation data^16^; no published chromatin accessibility data are available for comparison at these stages. In summary, we observe that all three molecular layers contain information to separate cells by stage and identity.

## Exit from pluripotency is associated with a sharp gain in DNA methylation and a gradual decrease in chromatin accessibility

To better understand epigenomic dynamics during development, we characterised the changes in DNA methylation and chromatin accessibility during each stage transition. CpG methylation levels rise from ~25% to ~75% in the embryonic tissue and ~50% in the extra-embryonic tissue (**Figure S9**), mainly driven by a *de novo* methylation wave from E4.5 to E5.5 that preferentially targets CpG-poor genomic loci^16,17,44^ (**Figure 1e,g,h**). Subsequent stage transitions induce relatively small global changes, but are instead associated with prominent local methylation processes, including X-chromosome inactivation in female embryos^17,45,46^ (**Figure S10**).

In contrast to the sharp *de novo* methylation dynamics between E4.5 and E5.5, we observed a more gradual decline in global chromatin accessibility from ~38% at E4.5 to ~30% at E7.5, with no differences between embryonic and extraembryonic tissues (**Figure S9**). However, consistent with the DNA methylation changes, CpG rich regions remain more accessible than CpG poor regions of the genome (**Figure 1f,g,h**)

To relate the epigenetic changes to transcriptional dynamics, we calculated, for each gene, the correlation coefficient across all cells between their RNA expression and the corresponding DNA methylation or chromatin accessibility levels at their promoters. Out of 5,000 genes tested (see **Methods**) we identified 205 genes whose expression shows significant correlation with promoter DNA methylation and 96 that show a significant correlation with chromatin accessibility (FDR<10%; **Figure 2a-b**).

Inspection of the dynamics for these associated loci reveals the repression of pluripotency and germ cell markers, including *Dppa4, Dppa5a, Zfp42, Tex19.1* and *Pou3f1* (**Figure 2a,c**), largely reflecting the genome-wide trend of DNA methylation gain and chromatin closure (**Figure 2c**). More globally, 41% and 32% of the genes that are downregulated after E4.5 show significant negative associations between RNA expression and promoter methylation and positive associations with chromatin accessibility, respectively. In contrast, only 9% and 1% of the genes found upregulated at E7.5 show a significant correlation between RNA expression and promoter methylation or accessibility, respectively (**Figure 2b, Figure S11**). This suggests that the upregulation of these genes is more likely controlled by other regulatory elements; a hypothesis that is further explored below.

In summary, we show that postimplantation development is characterised by increased DNA methylation and the global reduction in chromatin accessibility, which are associated with repression of pluripotency. Notably, this is opposite to the dynamics of preimplantation development, which has been found to be associated with an increase in chromatin accessibility and the removal of DNA methylation thereby establishing the naive pluripotent state^14,15^.

## Multi-omics factor analysis reveals connections between the transcriptome and the epigenetic state in enhancers during germ layer formation

Following the characterisation of the epigenetic landscape associated with the exit from pluripotency, we next explored associations of all three molecular layers with germ layer commitment at E7.5. To do this, we employed Multi-Omics Factor Analysis (MOFA)^47^, a method that performs unsupervised dimensionality reduction simultaneously across multiple data modalities, thereby capturing the global sources of cell-to-cell variability via a small number of inferred factors. Importantly, MOFA identifies whether factors are unique to a single data type or shared across multiple data types, thereby revealing the extent of covariation between different data modalities (**Figure S12**).

As input to the model we used the RNA expression profiles and the DNA methylation and chromatin accessibility data quantified over putative regulatory elements. Our observation that epigenetic changes in promoters show little association with genes upregulated at E7.5, prompted us to consider an extended set of regulatory elements. Specifically, we used histone ChIP-seq profiles specific to each of the three germ layers (Xiang Y. and Xie W., manuscript in preparation) to define enhancer elements (distal H3K27ac sites^48,49^) and active chromatin including transcription start sites (H3K4me3^50^), (**Figure S13**). Based on these annotations, we consider lineage-specific methylation and accessibility profiles as input data modalities for MOFA.

MOFA identified 7 robust factors (**Methods, Figure 3a**) with the first two (Factor 1 and Factor 2, sorted by variance explained) capturing the variation associated with the formation of the three germ layers (**Figure 3b**). Notably, for these two factors, MOFA links variation at the RNA level to concerted DNA methylation and chromatin accessibility changes at lineage-specific enhancer marks (**Figure 3a, Figure S14, S15**). In contrast, epigenetic changes at promoters show much weaker associations with the two lineaging factors than those at enhancer marks (**Figure 3a Figure S16,S17**). This supports observations in studies in other species that identified distal elements as lineage-driving regulatory regions^26,31^. Inspection of gene-enhancer associations identified enhancers linked to key germ layers including *Snai1, Lefty2* and *Mesp1* (mesoderm), *Sox17, Krt18* (endoderm), and *Pou3f1,Nav2* (ectoderm) (**Figure 3c**). Intriguingly, ectoderm-specific enhancers are less associated with lineage diversity than their meso- and endoderm counterparts (**Figure 3a, Figure S17**), suggesting an asymmetric contribution of epigenetic modifications to germ layer commitment. This finding is explored further below.

The five remaining factors correspond to transcriptional signatures that reflect the underlying spatial distribution of cells, such as anterior-posterior axial patterning (Factor 4, **Figure S18**); cell cycle (Factor 5, **Figure S19**); sublineaging events such as notochord formation (Factor 6, **Figure S20**), mesoderm patterning (Factor 7, **Figure S21**); and technical variability such as the cellular detection rate^51^ (Factor 3, **Figure S22**).

Finally, we sought to identify transcription factors (TF) that could drive or respond to the epigenetic changes in germ layer commitment. For each germ layer, we performed motif enrichment at enhancer sites that were differentially accessible between the germ layer of interest and the remaining two germ layers and intersected this with differential gene expression of the corresponding TF (**Methods, Figure S23**). Lineage-specific enhancers were enriched for binding sites associated with key developmental TFs, including POU3F1, SOX2 for ectoderm^52,53^; SOX17, FOXA2 for endoderm^54,55^; and GATA4, TWIST1 for mesoderm^56,57^ (**Figure 3d**).

## Mesoderm and endoderm enhancers undergo concerted demethylation and chromatin opening upon lineage specification

Following the characterisation of the molecular landscape of the three germ layers at E7.5, we next investigated the dynamics of establishment of methylation and accessibility states at lineage-defining enhancers (**Figure 4a and Figure S24**).

Endoderm and mesoderm-specific enhancers initially undergo a dramatic increase in DNA methylation from an average of ~25% to ~75% in all cell types, consistent with the genome-wide dynamics (**Figure S24-25**). Upon lineage specification, they undergo concerted demethylation to ~50% in a cell type specific manner, leading to the establishment of meso- and endoderm epigenetic profiles. The opposite pattern is observed for chromatin accessibility; the meso- and endoderm enhancer regions initially decrease from ~40% to ~30% accessibility, consistent again with the genome-wide dynamics (**Figure S24-25**), before becoming more accessible (~45%) upon lineage specification. The general dynamics of demethylation and chromatin opening of enhancers during embryogenesis seem thus to be conserved in zebrafish, Xenopus, and mouse^58^.

## Ectoderm enhancers are primed in the early epiblast supporting a default model of ectoderm commitment *in vivo*

In striking contrast to the mesoderm and endoderm enhancers, the epigenetic profiles in ectoderm enhancers are established as early as the E4.5 epiblast, and are protected from the global repressive dynamics, remaining accessible and hypomethylated until E7.5 in the ectoderm (**Figure 4a and Figure S24-25**). Only in cells committed (on the basis of their RNA profile) to a meso- or an endodermal state, are the ectoderm enhancers partially repressed (**Figure 4a**). Consistently, when measuring the accessibility dynamics at sites containing sequence motifs for SOX2, an ectoderm-defining TF, we find that these motifs are already accessible in the epiblast and lose accessibility specifically upon mesoderm and endoderm commitment (**Figure 4b**). Conversely, motifs associated with endoderm and mesoderm-defining TFs (FOXA2 and TWIST, respectively) have low accessibility in the epiblast and only become accessible in their respective lineages at E7.5.

Finally, to determine whether germ layer specific enhancers follow the same epigenetic dynamics *in vitro* we overlapped their genomic locations with ChIP-seq data for H3K27ac in mouse Epiblast Stem Cells (mEpiSC)^59^. Consistently, mEpiSCs show an ectoderm-like enhancer landscape with ~30% of ectoderm enhancers overlapping with H3K27ac marked regions in mEpiSC, compared to ~12% for mesoderm and endoderm enhancers (**Figure S26**). Additionally, as in *in vivo* epiblast cells, we find that chromatin accessibility and DNA methylation profiles of ectoderm enhancers are also found to be accessible and unmethylated in mouse Embryonic Stem Cells (ESCs) (**Figure S27**). This is consistent with a recent study describing enhancer priming for some lineage specific genes in ESCs^60^. Thus, ectoderm enhancers are epigenetically primed both *in vivo* and *in vitro*, while meso- and endoderm enhancers only acquire their unique epigenetic state at the time of germ layer specification.

**Figure 1:**
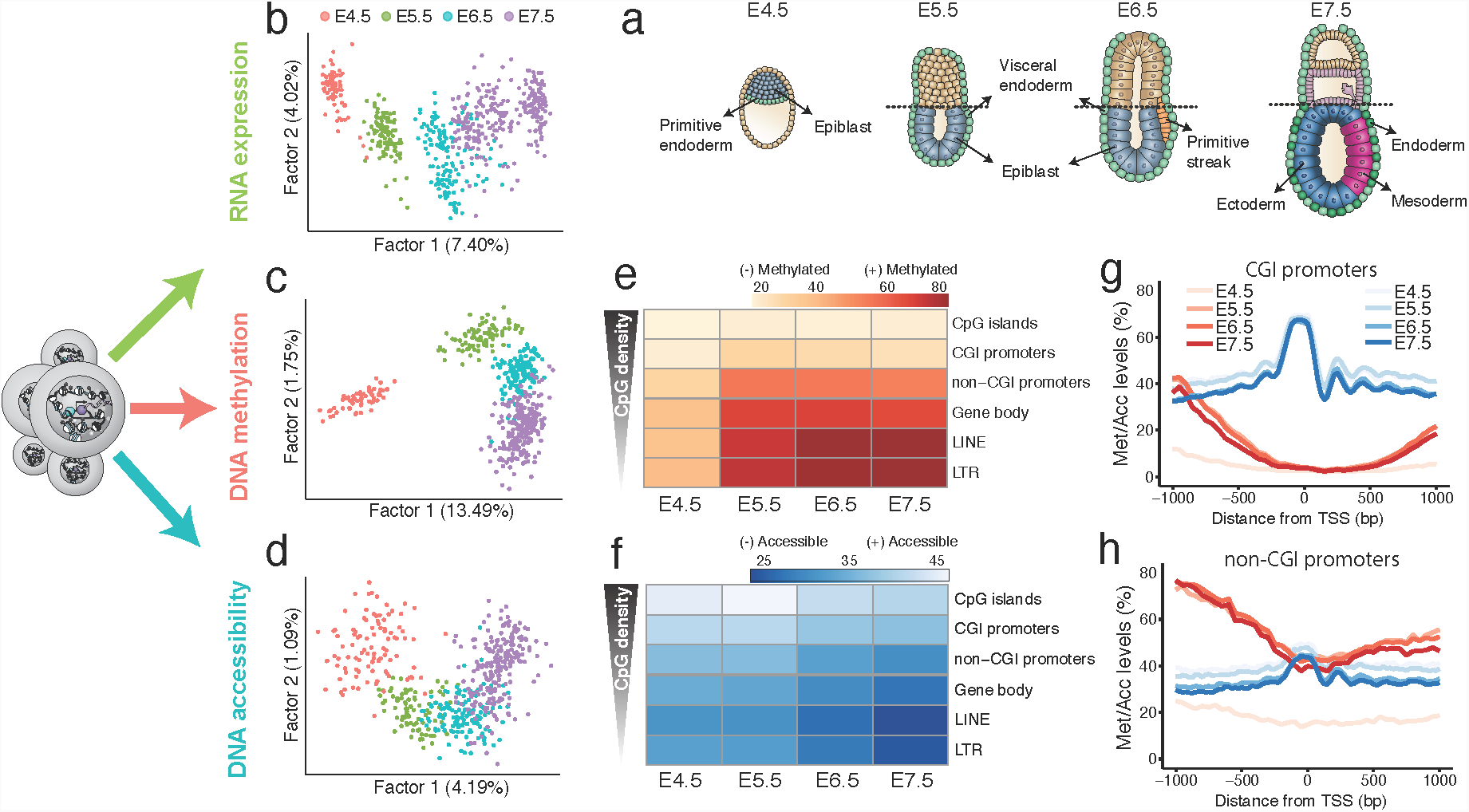
Single cell triple-omics profiling of mouse gastrulation. (a)Schematic of the developing mouse embryo, with stages and lineages analysed annotated. (b)Factor analysis projection of RNA expression data. (c)Factor analysis projection of DNA methylation data, quantified using non-overlapping 5kb windows along the genome. (d)Factor analysis projection of chromatin accessibility data, quantified using non-overlapping 100bp windows along the genome. All three molecular layers distinguish developmental stage. (e)Heatmap of DNA methylation levels (%) per stage and genomic context. (f)Heatmap of chromatin accessibility levels (%) per stage and genomic context. (g,h) DNA methylation (red) and chromatin accessibility (blue) profiles of (g) CGI promoters and (h) non-CGI promoters. Shown are genome-wide profiles, calculated using running averages of DNA methylation (red) and the chromatin accessibility (blue) (consecutive non-overlapping 50bp windows) across all cells within a stage.

**Figure 2:**
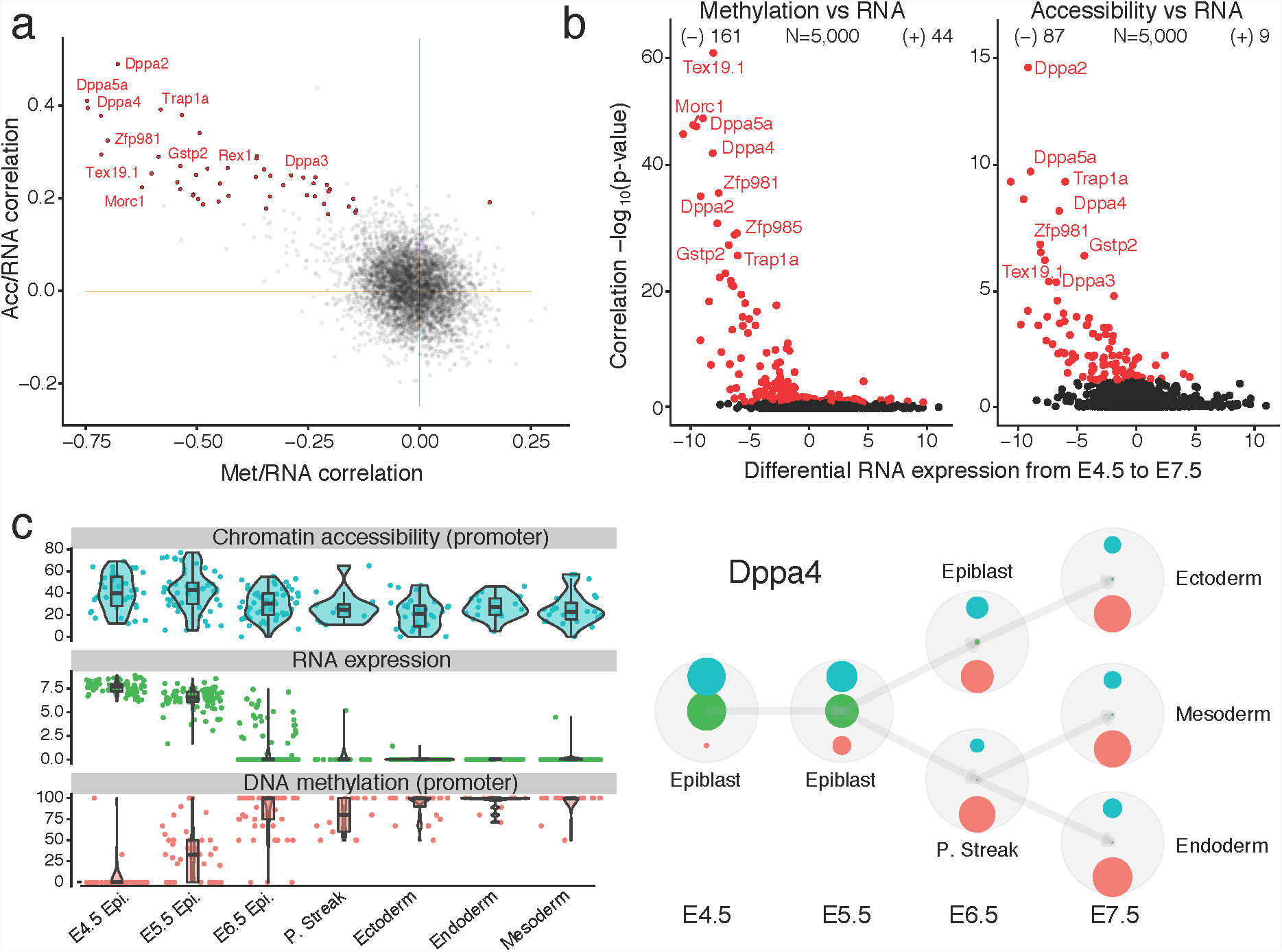
Epigenetic restriction in promoters is associated with the repression of pluripotency markers. (a)Scatter plot of correlation coefficients between promoter methylation and RNA expression (Met/RNA, x-axis), and promoter accessibility and RNA expression (Acc/RNA, y-axis). As in (b), each dot corresponds to one gene. Significant associations for both correlation types (FDR*<*10%) are coloured in red. Labels denote known pluripotency markers. (b)Volcano plots display differential RNA expression levels between E4.5 and E7.5 cells (difference in mean log2 counts, x-axis) versus adjusted correlation p-values (FDR*<*10%, Benjamini-Hochberg correction), for Met/RNA (left) and Acc/RNA (right). Negative values for differential RNA expression indicate genes are more expressed in E4.5, whereas positive values indicate genes that are more expressed in E7.5. (c)Illustrative example of epigenetic repression of the pluripotency associated gene Dppa4. Box and violin plots (left) show the distribution of chromatin accessibility (% levels, blue), RNA expression (log2 counts, green) and DNA methylation (% levels, red) values per stage and lineage. Box plots show median coverage and the first and third quartile, whiskers show 1.5xthe interquartile range. Each dot corresponds to one cell. Bubble plots (right) depict the relative values of chromatin accessibility (blue), RNA expression (green) and DNA methylation (red) at each stage and lineage. The size of each embedded circle represents a scaled value of each molecular layer (see Methods). Arrows depict known lineage transitions.

**Figure 3:**
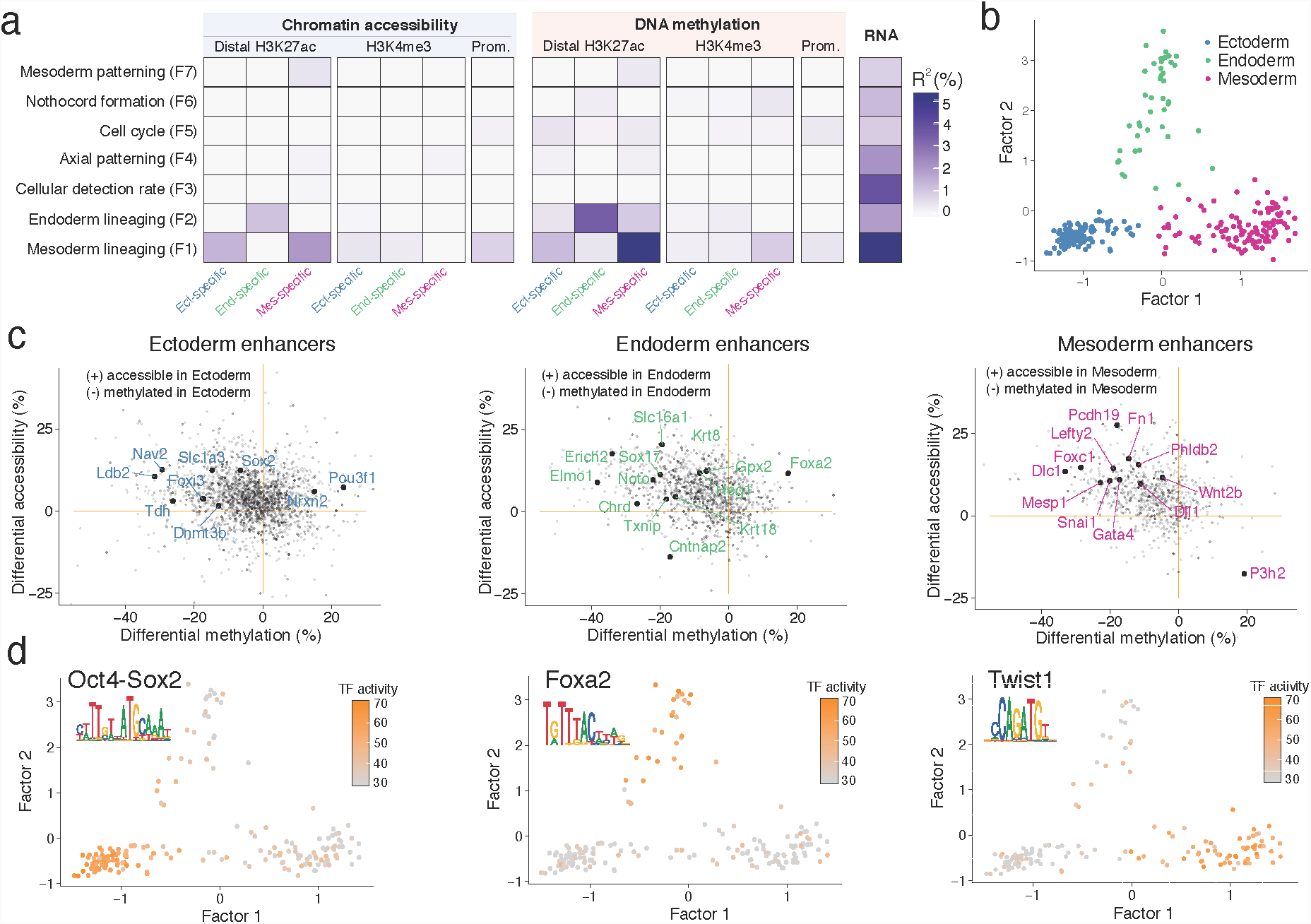
Multi-omics factor analysis reveals lineage-specific connections between transcriptome and epigenome variation at enhancer elements. (a)Percentage of variance explained (*R*^2^) by each MOFA factor (rows) across data modalities (columns). Data modalities were split by assay (red=methylation, blue=accessibility, green=RNA) and genomic context (promoters and lineage-specific H3K4me3 and distal H3K27ac ChIP-seq peaks). Factors are ordered by the average fraction of variance explained across all data modalities and annotated based on the inspection of the feature loadings and the visualisation of the latent space (see Figure S12). (b)Scatter plot showing the cells in a low dimensional representation using MOFA Factor 1 (x-axis) and MOFA Factor 2 (y-axis). Colours denote germ layer, assigned to each cell using the RNA profile (see Figure S4). (c)Scatter plots showing differential DNA methylation (%, x-axis) and chromatin accessibility (%, y-axis) at distal H3K27ac sites (enhancers). Shown are ectoderm vs non-ectoderm cells (left), endoderm vs non-endoderm cells (middle) and mesoderm vs non-mesoderm cells (right). Labeled big dots depict significant gene-enhancer pairs (FDR*<*10%) that also show significant lineage-specific RNA expression. (d)Scatter plot showing the cells in a low dimensional representation using the first two MOFA Factors as in (b). The color depicts transcription factor activity at the corresponding enhancer sites (left for ectoderm, middle for endoderm and right for mesoderm). Transcription factor activity is defined as average accessibility (%) across all motif instances within the set of lineage-specific enhancers.

**Figure 4:**
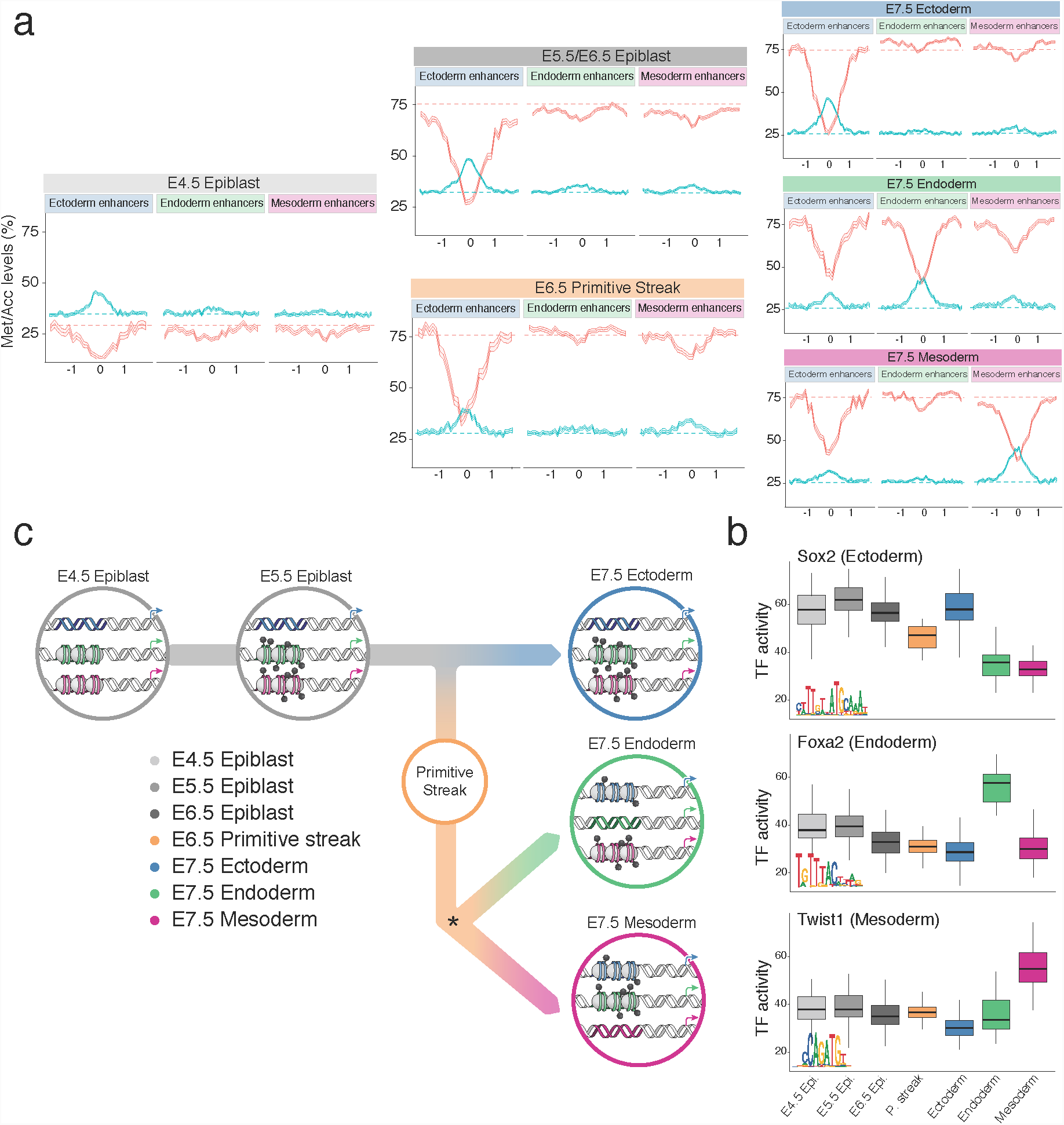
Epigenome dynamics of lineage-specific enhancers across development. (a)DNA methylation (red) and chromatin accessibility (blue) profiles of E7.5 lineage-specific enhancer sites at each stage and lineage. Shown are running averages of the DNA methylation (red) and the chromatin accessibility (blue) in consecutive non-overlapping 50bp windows. Solid line displays the mean across all cells (within a given lineage) and shading displays the corresponding standard deviation. E5.5 and E6.5 epiblast cells showed similar profiles and are combined. (b)Box plots showing the distribution of Transcription Factor (TF) activity at each stage and lineage. Each data point is a cell and TF activity is defined as average accessibility (%) across all motif instances within the set of lineage-specific enhancers in a given cell. (c)Illustration of the enhancer epigenetic dynamics during the hierarchical establishment of the epigenetic profiles associated with each germ layer.

## Conclusions

Our results show that even though pluripotent epiblast cells are clearly distinct from the E7.5 ectoderm at the transcriptional level (**Figure S28**), they are already epigenetically primed at ectoderm-defining enhancer sites. Hence, although lineage specific epigenetic marks at enhancers delineate each germ layer at E7.5 in a similar fashion, how these marks are established during development differs substantially between the ectoderm, and the mesoderm and endoderm. This finding supports the existence of a ‘default’ path in the Waddington landscape, providing a potential mechanism for the phenomenon of ‘default’ differentiation of neurectodermal tissue from ESCs^61–63^. Endoderm and mesoderm are thus actively diverted from the default path by signalling in the primitive streak which presumably induces demethylation and chromatin opening at the corresponding enhancers elements^26,32,64,65^. Hence, at least during gastrulation, lineages are defined by a hierarchical, or asymmetric, epigenetic model (**Figure 4c)**.

More generally, our discovery has important implications for the role of the epigenome in defining lineage commitment. Our work, and other recent studies also suggest chromatin priming may in certain circumstances precede overt cell fate decisions^26,60,66^. Additionally, we speculate that asymmetric epigenetic priming, where cells are epigenetically primed for a default cell type, may be a more general and poorly understood feature of differentiation and lineage commitment *in vivo*. Future studies that use multi-omics approaches to dissect cell populations at different stages of development therefore have the potential to transform our understanding of cell fate decision making, with important implications for stem cell biology and medicine.

## Methods

### Embryos and single cell isolation

All mice used in this study were C57BL/6Babr and were bred and maintained in the Babraham Institute Biological Support Unit. Ambient temperature was ~19-21°C and relative humidity 52%. Lighting was provided on a 12 hour light: 12 hour dark cycle including 15 min ‘dawn’ and ‘dusk’ periods of subdued lighting. After weaning, mice were transferred to individually ventilated cages with 1-5 mice per cage. Mice were fed CRM (P) VP diet (Special Diet Services) *ad libitum* and received seeds (e.g. sunflower, millet) at the time of cage-cleaning as part of their environmental enrichment. All mouse experimentation was approved by the Babraham Institute Animal Welfare and Ethical Review Body. Animal husbandry and experimentation complied with existing European Union and United Kingdom Home Office legislation and local standards. Single-cells from E4.5 to E6.5 embryos were collected as described^8^. E7.5 embryos were dissected to remove extra-embryonic tissue and dissociated in TryplE for 10 minutes at room temperature. Undigested portions were physically removed and the remainder filtered through a 30 μm filter prior to isolation using flow cytometry.

### scNMT-seq library preparation

Single-cells were sorted either manually (E4.5, E5.5) or using flow cytometry (E6.5, E7.5) into 96 well PCR plates containing 2.5μl of methylase reaction buffer (1× M.CviPI Reaction buffer (NEB), 2U M.CviPI (NEB), 160μM S-adenosylmethionine (NEB), 1Uμl^-1^ RNasein (Promega), 0.1% IGEPAL (Sigma)). Samples were incubated for 15 minutes at 37°C to methylate accessible chromatin before the reaction was stopped with the addition of RLT plus buffer (Qiagen) and samples frozen down and stored at -80°C prior to processing. Poly-A RNA was captured on oligo-dT conjugated to magnetic beads and amplified cDNA was prepared according to the G&T-seq^67^ and Smartseq2 protocols^68^. The lysate containing gDNA was purified on AMPureXP beads before bisulfite-seq libraries were prepared according to the scBS-seq protocol^69^.

### Sequencing

All sequencing was carried out on a NextSeq500 instrument in high-output mode using 75bp paired end reads. Libraries were pooled as 48-plexes (BS-seq) or 384-plexes (RNA-seq). This yielded a mean raw sequencing depth of 8.0 million (BS-seq) and 0.98 million (RNA-seq) paired-end reads per cell.

### RNA-seq alignment and quantification

RNA-seq libraries were aligned to the GRCm38 mouse genome build using HiSat2^70^ (v2.1.0) using options --dta --sp 1000,1000 --no-mixed --no-discordant, yielding a mean of 611,000 aligned reads per cell.

Subsequently, gene expression counts were quantified from the mapped reads using featureCounts^71^ with the Ensembl gene annotation^72^ (version 87). Only protein-coding genes matching canonical chromosomes were considered. The read counts were log-transformed and size-factor adjusted^73^.

### BS-seq alignment and methylation/accessibility quantification

BS-seq libraries were aligned to the bisuflite converted GRCm38 mouse genome using Bismark^74^ (v0.19.1) in single-end nondirectional mode. Following the removal of PCR duplicates, we retained a mean of 1.6 million reads per cell. Methylation calling and separation of endogenous methylation (from A-C-G and T-C-G trinucleotides) and chromatin accessibility (G-C-A, G-C-C and G-C-T trinucleotides) was performed with Bismark using the --NOMe option of the coverage2cytosine script.

Following our previous approach^75^, individual CpG or GpC sites in each cell were modelled using a binomial distribution where the number of successes is the number of reads that support methylation and the number of trials is the total number of reads. A CpG methylation or GpC accessibility rate for each site and cell was calculated by maximum likelihood. The rates were subsequently rounded to the nearest integer (0 or 1).

When aggregating over genomic features, CpG methylation and GpC accessibility rates were computed assuming a binomial model, with the number of trials being the number of sites and the number of successes being the number of methylated sites.

### ChIP-seq experiments

Three germ layers were collected as described previously^76^. Briefly, embryos were dissected from uterus and decidua. Reichert’s membrane was further removed using syringe needles. Embryonic regions were separated from extraembryonic tissues and incubated in pancreatic and trypsin enzyme solution for 5 minutes. After collecting endoderm by gently sucking the embryonic part into a capillary pipet, glass needles were used to cut off both mesoderm wings, which is adjacent to the primitive streak region. Finally, ectoderm was collected by removing posterior primitive streak.

STAR ChIP-seq of histone modifications was performed as described previously^77^. Briefly, three germ layers were lysed and fragmented by MNase. The histone and DNA solution was incubated with 1μg H3K27ac (Active motif, 39133) or H3K4me3 (homemade^77^) overnight. After eluting DNA from histone and repairing dephosphorylate 3’ end, the resulting DNA was subjected to TELP library preparation as described^78^.

### ChIP-seq data processing

H3K4me3 and H3K27ac ChIP-seq libraries produced from two biological replicates were sequenced as single-end or paired-end runs with read lengths of 50, 100 or 125 bp (see **Table S1** for details). Read 2 was excluded from the analysis for paired end samples because of low quality scores. Reads were trimmed using Trim Galore (v0.4.5, cutadapt 1.15, single end mode) and mapped to *M. musculus* GRCm38 using Bowtie2^79^ (v2.3.2). All analyses were performed using SeqMonk (www.bioinformatics.babraham.ac.uk/projects/seqmonk/). For quantitation, read length was extended to 300 bp and regions of coverage outliers and extreme strand bias excluded. Comparison of data sets with different read lengths did not reveal major mapping differences, and thus, mapped, extended reads were merged for samples that were sequenced across more than one lane. Samples were overall similar regarding total mapped read numbers, distribution of reads and ChIP enrichment.

To best represent the underlying ChIP-seq signal, different methods to define enriched genomic regions were used for H3K4me3 and H3K27ac marks. For H3K4me3, a SeqMonk implementation of MACS^80^ with the local rescoring step omitted was used (p<10-15, fragment size 300 bp), and enriched regions closer than 100 bp were merged. Peaks were called separately for each lineage. For H3K27ac, reads were quantitated per 500 bp tiles correcting per million total reads and excluding duplicate reads. Smoothing subtraction quantitation was used to identify local maxima (value > 1), and peaks closer than 500 bp apart were merged. Lineage specific peak annotations exclude peaks that are also present in one of the other lineages, and only peaks present in both replicates were considered (**Figure S13**).

### scRNA-seq quality control

Cells with less than 100,000 mapped reads and with less than 2,500 expressed genes were excluded. In total, 685 cells passed quality control on the basis of their mRNA expression levels (**Figure S1**).

### scBS-seq quality control

Cells with less than 100,000 CpG sites and 1,000,000 GpC sites covered were discarded. In total, 597 cells passed quality control for DNA methylation and 576 cells passed quality control for chromatin accessibility (**Figure S1**).

### Lineage assignment

At each developmental stage, lineages were assigned using SC3^81^ based upon the RNA expression profiles (**Figure S2-5**). For the assignment of the E6.5 primitive streak cells, a Principal Component Analysis was performed using selected gene markers. Cells were classified as epiblast or primitive streak based on a selected threshold on the first component (**Figure S4**).

### Correlation analysis

To identify genes with an association between the mRNA expression and promoter epigenetic status, we calculated, for each gene, the correlation coefficient across all cells between its RNA expression and the corresponding DNA methylation or chromatin accessibility levels at the gene’s promoter (+/-2kb around transcription start site).

As a filtering criterion, we required, for each genomic feature, a minimum number of 1 CpG (methylation) or 3 GpC (accessibility) measurements in at least 25 cells. Additionally, the top 5,000 most variable genes (across all cells) were selected, according to the rationale of independent filtering^82^. Two-tailed Student’s t-tests were performed to test for evidence against the null hypothesis of no correlation, and p-values were adjusted for multiple testing using the Benjamini–Hochberg procedure^83^.

### Differential DNA methylation and chromatin accessibility analysis

Differential analysis of DNA methylation and chromatin accessibility between different pre-defined groups was performed using a Fisher exact test independently for each genomic element

More precisely, cells were aggregated into two exclusive groups, defined by either stage or lineage, depending on the analysis. Then, for a given genomic element, we created a contingency table by aggregating (across cells) the number of methylated and unmethylated nucleotides in each group as follows:

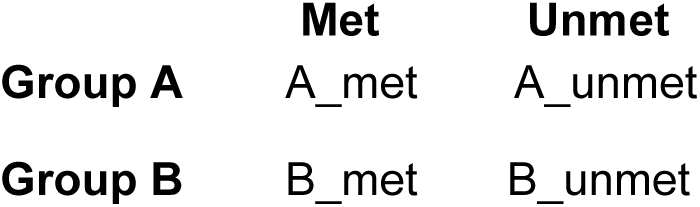

Multiple testing correction was applied using the Benjamini-Hochberg procedure. As a filtering criteria, we required 1 CpG (methylation) and 3 GpC (accessibility) observations in at least 5 cells per group. Non-variable regions were filtered out prior to differential testing.

### Motif enrichment

To find transcription factor motifs enriched in lineage-associated sites, we used lineage-specific H3K27ac sites that were identified as differentially accessible between lineages as explained above. We tested for enrichment over a background of all H3K27ac sites. Fasta files were generated using the Bioconductor package *BSgenome*^*84*^ and enrichment was tested using ame (meme suite^85^ v4.10.1) with parameters *--method fisher--scoring avg*. Position frequency matrices were downloaded from the Jaspar core vertebrates database^86^.

### Differential RNA expression analysis

Differential RNA expression analysis between pre-specified groups of interest was performed using the likelihood ratio test from edgeR^87^. Significant hits were called with a 1% False Discovery Rate (Benjamini-Hochberg procedure) and a minimum log2 fold change of 1.Genes with non-variable expression were filtered out prior to differential testing^82^.

### Additional data sets

For published datasets, we downloaded fastq files from GEO (GSE76505^16^) and used the GRCm38 version of the mouse genome for all alignments. RNA-seq libraries were processed using Hisat2^70^ (v2.1.0). Gene expression counts were quantified from the mapped reads using featureCounts^71^ with the Ensembl gene annotation^72^ (version 87). BS-TELP^16^ libraries were processed using Bismark^74^ (v0.19.0) with default parameters. Methylation rates were computed at predefined genomic annotations as the fraction of methylated CpG sites divided by the sum of all CpG sites within the given region.

### Dimensionality reduction using Factor Analysis

To show that stages and lineages have different DNA methylation and chromatin accessibility profiles we performed dimensionality reduction using unsupervised running windows along the genome. To handle the large amount of missing values and to estimate the variance, we used MOFA^47^. To ensure the data followed a Gaussian distribution we calculated M-values from the beta values (rates).

Due to differences in genomic coverage^34^ we used a window size of 5kb for DNA methylation and a window size of 100bp for chromatin accessibility. To select informative sites and increase computational efficiency in the accessibility data, we first selected accessible (>75%) 100bp windows in pseudobulk E4.5 data (~250,000 sites), which is similar to approaches used to analyse single-cell ATAC-seq data^26^. For both data modalities, we fitted the model using the top 5,000 most variable sites. For every model, we learnt two latent factors, which were subsequently used for visualisation (**Figure 1a-c**, **Figure SF7-8**). Variance explained estimates were computed using the coefficient of determination as described in ^47^.

### Multi-Omics Factor Analysis (MOFA)

The input to MOFA is a set of matrices with dimensions (features, cells), defined as views. To quantify the contributions from all three molecular layers to cellular diversity during germ layer formation (**Figure 3**), we selected distal H3K27ac sites (enhancers) and H3K4me3 (active transcription start sites) as regulatory genomic contexts. Both annotations were defined using independently generated lineage-specific ChIP-seq data at E7.5. Additionally, we included promoters defined as 2kb windows upstream and downstream of a gene’s TSS. For each genomic context, we created a DNA methylation matrix and a chromatin accessibility matrix by quantifying M-values for each cell and feature (i.e. promoter or ChIP-seq peak). To reduce computational complexity, the top 2,500 most variable features across E7.5 cells were selected per view. For RNA expression, log normalised counts were quantified per gene and cell, and the top 2,500 most variable genes were selected. The same number of features was selected for each view to avoid class imbalance problems.

As a filtering criteria, genomic features were required to have a minimum of 1 CpG (methylation) or 3 GpC (accessibility) observed in at least 15 cells. Genes were required to have a minimum cellular detection rate of 25%. As MOFA handles large amounts of missing values, including cells missing a particular view^47^, no imputation was required.

Since parameters are randomly initialised, the optimisation procedure of MOFA is not guaranteed to find a consistent optimal solution on every single run, meaning that factors can vary between different model instances. To address this, we adopted the pipeline proposed in ^47^, we trained 10 model instances under different random initialisations, and we performed model selections based on the maximal Evidence Lower Bound (ELBO) value. The consistency of factors was assessed by calculating the Pearson correlation coefficients between every pair of factors across the 10 trials. All factors were found in at least 5 model instances and hence were considered robust for downstream analysis.

MOFA identifies the numbers of factors based on a minimum variance criterion. In this analysis, we used a minimum of 1% explained variance on the RNA expression view.

The downstream characterisation of the model output included several analyses:

- **Variance decomposition by factor:** The first step, after a model has been trained, is to disentangle the variation explained by each factor in each view. To this end, we compute the fraction of the variance explained (R^2^) by factor *k* in view *m* using the coefficient of determination^47^.
- **Visualisation of loadings:** For each factor, the model learns a loading per feature in each view. The absolute value of the loading can be interpreted as the strength of association between the observed values for the given feature and the ordination of samples defined by the factor. Hence, features with large loadings are highly correlated with the factor values, and thus inspecting the top features can give insights into the biological process underlying the heterogeneity captured by a latent factor.
- **Visualisation of factors:** each factor captures an independent dimension of heterogeneity. Mathematically, this translates into ordering samples in a real-valued latent space. Hence, similar to Principal Component Analysis, we plot factor values against each other to reveal the hidden structure of the data. For more details, see ^47^.
- **Feature set enrichment analysis**: when inspecting the loadings for a given factor, multiple features can be combined into a gene set-based annotation. For a given gene set G, we evaluate its significance via a parametric t-test, where we compare the weights of the foreground set (features that belong to the set G) and the background set (the weights of features that do not belong to the set G).

## Code availability

All analysis code is available at https://github.com/PMBio/gastrulation

## Data availability

Raw sequencing data together with processed files (RNA counts, CpG methylation reports, GpC accessibility reports) are available in the Gene Expression Omnibus under accession GSEXXX. All other data are available from the authors upon reasonable request.

## Author contributions

H.M. and W.R. conceived the project.

H.M., S.J.C., S.S., P.R-G., W.D., J.N. performed experiments.

R.A., S.J.C., F.B., L.C.S., X.I., C.A.K. and C.K. performed computational analysis.

F.K. processed and managed sequencing data.

Y.X. and C.H. performed ChIP-seq experiments.

R.A. S.J.C., L.C.S., W.D., O.S., J.C.M., W.R. interpreted results and drafted the manuscript. G.S., P.R-G., W.X. G.K., O.S., J.C.M., W.R. supervised the project

## Acknowledgements

R.A. is a member of Robinson College at University of Cambridge. We thank K. Tabbada and C. Murnane of the Babraham Next Generation Sequencing Facility for assistance with Illumina sequencing, members of of the Babraham Flow Cytometry Core Facility for cell sorting and the Babraham Biological Support Unit for animal work. We also thank Yu Zhang for help in processing the ChIP-seq data. L.C.S. is supported by EMBO postdoctoral fellowship (ALTF 417-2018). J.C.M. is supported by core funding from EMBL and CRUK. R.A. is supported by the EMBL International Predoc Programme. X.I.S. is supported by Wellcome Trust Grant 108438/E/15/Z, ‘New genetic, imaging and microfluidics technologies for single cell genomics’

## References

1. SolnicaKrezel, L. & Sepich, D. S. Gastrulation: making and shaping germ layers. Annu. Rev. Cell Dev. Biol. 28, 687–717 (2012).

2. Tam, P. P.L. & Loebel, D. A. F. Gene function in mouse embryogenesis: get set for gastrulation. Nat. Rev. Genet. 8, 368–381 (2007).

3. Griffiths, J. A., Scialdone, A. & Marioni, J. C. Using single cell genomics to understand developmental processes and cell fate decisions. Mol. Syst. Biol. 14, e8046 (2018).

4. Hadjantonakis, A. K. & Arias, A. M. Single Cell Approaches: Pandora’s Box of Developmental Mechanisms. Dev. Cell 38, 574–578 (2016).

5. Karaiskos, N. et al. The Drosophila embryo at single cell transcriptome resolution. Science 358, 194–199 (2017).

6. Arnold, S. J. & Robertson, E. J. Making a commitment: cell lineage allocation and axis patterning in the early mouse embryo. Nat. Rev. Mol. Cell Biol. 10, 91–103 (2009).

7. Scialdone, A. et al. Resolving early mesoderm diversification through single cell expression profiling. Nature 535, 289–293 (2016).

8. Mohammed, H. et al. Single Cell Landscape of Transcriptional Heterogeneity and Cell Fate Decisions during Mouse Early Gastrulation. Cell Rep. 20, 1215–1228 (2017).

9. Wen, J. et al. Single cell analysis reveals lineage segregation in early post implantation mouse embryos. J. Biol. Chem. 292, 9840–9854 (2017).

10. Ibarra­Soria, X. et al. Defining murine organogenesis at single cell resolution reveals a role for the leukotriene pathway in regulating blood progenitor formation. Nat. Cell Biol. 20, 127–134 (2018).

11. Peng, G. et al. Spatial Transcriptome for the Molecular Annotation of Lineage Fates and Cell Identity in Mid gastrula Mouse Embryo. Dev. Cell 36, 681–697 (2016).

12. Chan, M. et al. Molecular recording of mammalian embryogenesis. bioRxiv 384925 (2018). doi:10.1101/384925

13. Kelsey, G., Stegle, O. & Reik, W. Single cell epigenomics: Recording the past and predicting the future. Science 358, 69–75 (2017).

14. Wu, J. et al. The landscape of accessible chromatin in mammalian preimplantation embryos. Nature 534, 652–657 (2016).

15. Lu, F. et al. Establishing Chromatin Regulatory Landscape during Mouse Preimplantation Development. Cell 165, 1375–1388 (2016).

16. Zhang, Y. et al. Dynamic epigenomic landscapes during early lineage specification in mouse embryos. Nat. Genet. 50, 96–105 (2018).

17. Auclair, G., Guibert, S., Bender, A. & Weber, M. Ontogeny of CpG island methylation and specificity of DNMT3 methyltransferases during embryonic development in the mouse. Genome Biol. 15, 545 (2014).

18. Lee, H. J., Hore, T. A. & Reik, W. Reprogramming the methylome: erasing memory and creating diversity. Cell Stem Cell 14, 710–719 (2014).

19. Messerschmidt, D. M., Knowles, B. B. & Solter, D. DNA methylation dynamics during epigenetic reprogramming in the germline and preimplantation embryos. Genes Dev. 28, 812–828 (2014).

20. Smith, Z. D. et al. DNA methylation dynamics of the human preimplantation embryo. Nature 511, 611–615 (2014).

21. Smith, Z. D. et al. Epigenetic restriction of extraembryonic lineages mirrors the somatic transition to cancer. Nature 549, 543–547 (2017).

22. Guo, H. et al. The DNA methylation landscape of human early embryos. Nature 511, 606–610 (2014).

23. Smith, Z. D. & Meissner, A. DNA methylation: roles in mammalian development. Nat. Rev. Genet. 14, 204–220 (2013).

24. Zhu, P. et al. Single cell DNA methylome sequencing of human preimplantation embryos. Nat. Genet. 50, 12–19 (2018).

25. Xu, Q. & Xie, W. Epigenome in Early Mammalian Development: Inheritance, Reprogramming and Establishment. Trends Cell Biol. 28, 237–253 (2018).

26. Cusanovich, D. A. et al. The cis regulatory dynamics of embryonic development at single cell resolution. Nature 555, 538–542 (2018).

27. Cusanovich, D. A. et al. A Single Cell Atlas of In Vivo Mammalian Chromatin Accessibility. Cell 174, 1309–1324.e18 (2018).

28. McKay, D. J. & Lieb, J. D. A common set of DNA regulatory elements shapes Drosophila appendages. Dev. Cell 27, 306–318 (2013).

29. Lee, H. J. et al. Developmental enhancers revealed by extensive DNA methylome maps of zebrafish early embryos. Nat. Commun. 6, 6315 (2015).

30. Wiench, M. et al. DNA methylation status predicts cell type specific enhancer activity. EMBO J. 30, 3028–3039 (2011).

31. Daugherty, A. C. et al. Chromatin accessibility dynamics reveal novel functional enhancers in C. elegans. Genome Res. 27, 2096–2107 (2017).

32. Dai, H. Q. et al. TET mediated DNA demethylation controls gastrulation by regulating Lefty Nodal signalling. Nature 538, 528–532 (2016).

33. Okano, M., Bell, D. W., Haber, D. A. & Li, E. DNA methyltransferases Dnmt3a and Dnmt3b are essential for de novo methylation and mammalian development. Cell 99, 247–257 (1999).

34. Clark, S. J. et al. scNMT seq enables joint profiling of chromatin accessibility DNA methylation and transcription in single cells. Nat. Commun. 9, 781 (2018).

35. Angermueller, C. et al. Parallel single cell sequencing links transcriptional and epigenetic heterogeneity. Nat. Methods 13, 229–232 (2016).

36. Macaulay, I. C., Ponting, C. P. & Voet, T. Single Cell Multiomics: Multiple Measurements from Single Cells. Trends Genet. 33, 155–168 (2017).

37. Macaulay, I. C. et al. G&T seq: parallel sequencing of single cell genomes and transcriptomes. Nat. Methods 12, 519–522 (2015).

38. Dey, S. S., Kester, L., Spanjaard, B., Bienko, M. & van Oudenaarden, A. Integrated genome and transcriptome sequencing of the same cell. Nat. Biotechnol. 33, 285–289 (2015).

39. Guo, F. et al. Single cell multi omics sequencing of mouse early embryos and embryonic stem cells. Cell Res. 27, 967–988 (2017).

40. Pott, S. Simultaneous measurement of chromatin accessibility, DNA methylation, and nucleosome phasing in single cells. Elife 6, (2017).

41. Hu, Y. et al. Simultaneous profiling of transcriptome and DNA methylome from a single cell. Genome Biol. 17, 88 (2016).

42. Li, L. et al. Single cell multi omics sequencing of human early embryos. Nat. Cell Biol. 20, 847–858 (2018).

43. Cao, J. et al. Joint profiling of chromatin accessibility and gene expression in thousands of single cells. Science 361, 1380–1385 (2018).

44. Smith, Z. D. et al. A unique regulatory phase of DNA methylation in the early mammalian embryo. Nature 484, 339–344 (2012).

45. Gendrel, A. V. et al. Smchd1 dependent and independent pathways determine developmental dynamics of CpG island methylation on the inactive X chromosome. Dev. Cell 23, 265–279 (2012).

46. Norris, D. P., Brockdorff, N. & Rastan, S. Methylation status of CpG rich islands on active and inactive mouse X chromosomes. Mamm. Genome 1, 78–83 (1991).

47. Argelaguet, R. et al. Multi Omics Factor Analysis a framework for unsupervised integration of multi omics data sets. Mol. Syst. Biol. 14, e8124 (2018).

48. Creyghton, M. P. et al. Histone H3K27ac separates active from poised enhancers and predicts developmental state. Proc. Natl. Acad. Sci. U. S. A. 107, 21931–21936 (2010).

49. Calo, E. & Wysocka, J. Modification of enhancer chromatin: what, how, and why? Mol. Cell 49, 825–837 (2013).

50. Liang, G. et al. Distinct localization of histone H3 acetylation and H3 K4 methylation to the transcription start sites in the human genome. Proc. Natl. Acad. Sci. U. S. A. 101, 7357–7362 (2004).

51. Finak, G. et al. MAST: a flexible statistical framework for assessing transcriptional changes and characterizing heterogeneity in single cell RNA sequencing data. Genome Biol. 16, 278 (2015).

52. Zhang, S. & Cui, W. Sox2, a key factor in the regulation of pluripotency and neural differentiation. World J. Stem Cells 6, 305–311 (2014).

53. Zhu, Q. et al. The transcription factor Pou3f1 promotes neural fate commitment via activation of neural lineage genes and inhibition of external signaling pathways. Elife 3, (2014).

54. Burtscher, I. & Lickert, H. Foxa2 regulates polarity and epithelialization in the endoderm germ layer of the mouse embryo. Development 136, 1029–1038 (2009).

55. Séguin, C. A., Draper, J. S., Nagy, A. & Rossant, J. Establishment of endoderm progenitors by SOX transcription factor expression in human embryonic stem cells. Cell Stem Cell 3, 182–195 (2008).

56. Bildsoe, H. et al. The mesenchymal architecture of the cranial mesoderm of mouse embryos is disrupted by the loss of Twist1 function. Dev. Biol. 374, 295–307 (2013).

57. Molkentin, J. D., Lin, Q., Duncan, S. A. & Olson, E. N. Requirement of the transcription factor GATA4 for heart tube formation and ventral morphogenesis. Genes Dev. 1, 1061–1072 (1997).

58. Bogdanović, O. et al. Active DNA demethylation at enhancers during the vertebrate phylotypic period. Nat. Genet. 48, 417–426 (2016).

59. Factor, D. C. et al. Epigenomic comparison reveals activation of ‘seed’ enhancers during transition from naive to primed pluripotency. Cell Stem Cell 14, 854–863 (2014).

60. Kim, H. S. et al. Pluripotency factors functionally premark cell type restricted enhancers in ES cells. Nature 556, 510–514 (2018).

61. Grunz, H. & Tacke, L. Neural differentiation of Xenopus Zueuis ectoderm takes place after disaggregation and delayed reaggregation without inducer. Cell Differ. Dev. 28, 211–218 (1989).

62. Tropepe, V. et al. Direct neural fate specification from embryonic stem cells: a primitive mammalian neural stem cell stage acquired through a default mechanism. Neuron 30, 65–78 (2001).

63. Muñoz Sanjuán, I. & Brivanlou, A. H. Neural induction, the default model and embryonic stem cells. Nat. Rev. Neurosci. 3, 271–280 (2002).

64. Sardina, J. L. et al. Transcription Factors Drive Tet2 Mediated Enhancer Demethylation to Reprogram Cell Fate. Cell Stem Cell (2018). doi:10.1016/j.stem.2018.08.016

65. Simon, C. S. et al. Functional characterisation of cis regulatory elements governing dynamic Eomes expression in the early mouse embryo. Development 144, 1249–1260 (2017).

66. Heinz, S. et al. Simple combinations of lineage determining transcription factors prime cis regulatory elements required for macrophage and B cell identities. Mol. Cell 38, 576–589 (2010).

67. Macaulay, I. C. et al. Separation and parallel sequencing of the genomes and transcriptomes of single cells using G&T seq. Nat. Protoc. 11, 2081–2103 (2016).

68. Picelli, S. et al. Full length RNA seq from single cells using Smart seq2. Nat. Protoc. 9, 171–181 (2014).

69. Clark, S. J. et al. Genome wide base resolution mapping of DNA methylation in single cells using single cell bisulfite sequencing (scBS seq). Nat. Protoc. 12, 534–547 (2017).

70. Kim, D., Langmead, B. & Salzberg, S. L. HISAT: a fast spliced aligner with low memory requirements. Nat. Methods 12, 357–360 (2015).

71. Liao, Y., Smyth, G. K. & Shi, W. featureCounts: an efficient general purpose program for assigning sequence reads to genomic features. Bioinformatics 30, 923–930 (2014).

72. Yates, A. et al. Ensembl 2016. Nucleic Acids Res. 44, D710–6 (2016).

73. Lun, A. T. L., McCarthy, D. J. & Marioni, J. C. A step by step workflow for low level analysis of single cell RNA seq data with Bioconductor. F1000Res. 5, 2122 (2016).

74. Krueger, F. & Andrews, S. R. Bismark: a flexible aligner and methylation caller for Bisulfite Seq applications. Bioinformatics 27, 1571–1572 (2011).

75. Smallwood, S. A. et al. Single cell genome wide bisulfite sequencing for assessing epigenetic heterogeneity. Nat. Methods 11, 817–820 (2014).

76. Nagy, A., Gertsenstein, M., Vintersten, K. & Behringer, R. Collecting zygotes and removing cumulus cells with hyaluronidase. CSH Protoc. 2006, (2006).

77. Zhang, B. et al. Allelic reprogramming of the histone modification H3K4me3 in early mammalian development. Nature 537, 553–557 (2016).

78. Peng, X. et al. TELP, a sensitive and versatile library construction method for next generation sequencing. Nucleic Acids Res. 43, e35 (2015).

79. Langmead, B. & Salzberg, S. L. Fast gapped read alignment with Bowtie 2. Nat. Methods 9, 357–359 (2012).

80. Zhang, Y. et al. Model based analysis of ChIP Seq (MACS). Genome Biol. 9, R137 (2008).

81. Kiselev, V. Y. et al. SC3: consensus clustering of single cell RNA seq data. Nat. Methods 14, 483–486 (2017).

82. Bourgon, R., Gentleman, R. & Huber, W. Independent filtering increases detection power for high throughput experiments. Proc. Natl. Acad. Sci. U. S. A. 107, 9546–9551 (2010).

83. Benjamini, Y. & Hochberg, Y. Controlling the False Discovery Rate: A Practical and Powerful Approach to Multiple Testing. J. R. Stat. Soc. Series B Stat. Methodol. 57, 289–300 (1995).

84. Pages, H. BSgenome: Infrastructure for Biostrings based genome data packages. R package version 1, (2012).

85. McLeay, R. C. & Bailey, T. L. Motif Enrichment Analysis: a unified framework and an evaluation on ChIP data. BMC Bioinformatics 11, 165 (2010).

86. Khan, A. et al. JASPAR 2018: update of the open access database of transcription factor binding profiles and its web framework. Nucleic Acids Res. 46, D260–D266 (2018).

87. Robinson, M. D., McCarthy, D. J. & Smyth, G. K. edgeR: a Bioconductor package for differential expression analysis of digital gene expression data. Bioinformatics 26, 139–140 (2010).

