## Supplementary Figures for "Single cell multi-omics profiling reveals a hierarchical epigenetic landscape during mammalian germ layer specification"

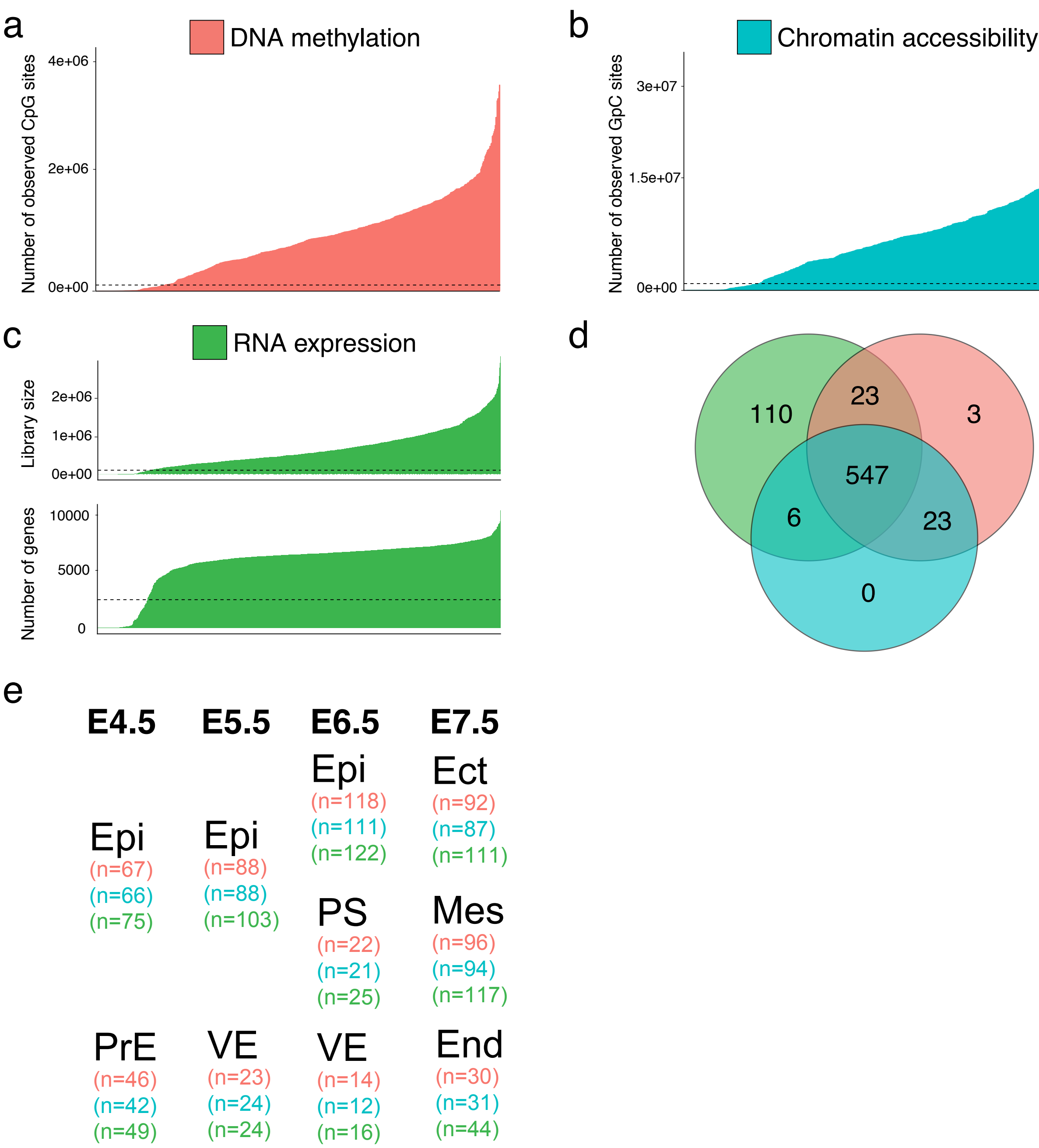


**Figure S1: scNMT-seq quality controls.**
(a-b) Number of observed cytosines in (a) CpG (red) or (b) GpC (blue) contexts respectively. Each bar corresponds to one cell. Cells below a set threshold (dotted lines) were discarded on the basis of poor quality.
(c) Library size (top, total number of reads) and number of expressed genes (bottom, log2 normalised read counts>0) per cell. Cells below a set threshold (dotted lines) were discarded on the basis of poor quality.
(d) Venn Diagram displaying the number of cells that passed quality control for RNA expression (green), DNA methylation (red), chromatin accessibility (blue) and overlapping (multi-omics) information.
(e) Number of cells that passed quality control for each individual omic, grouped by stage and lineage.

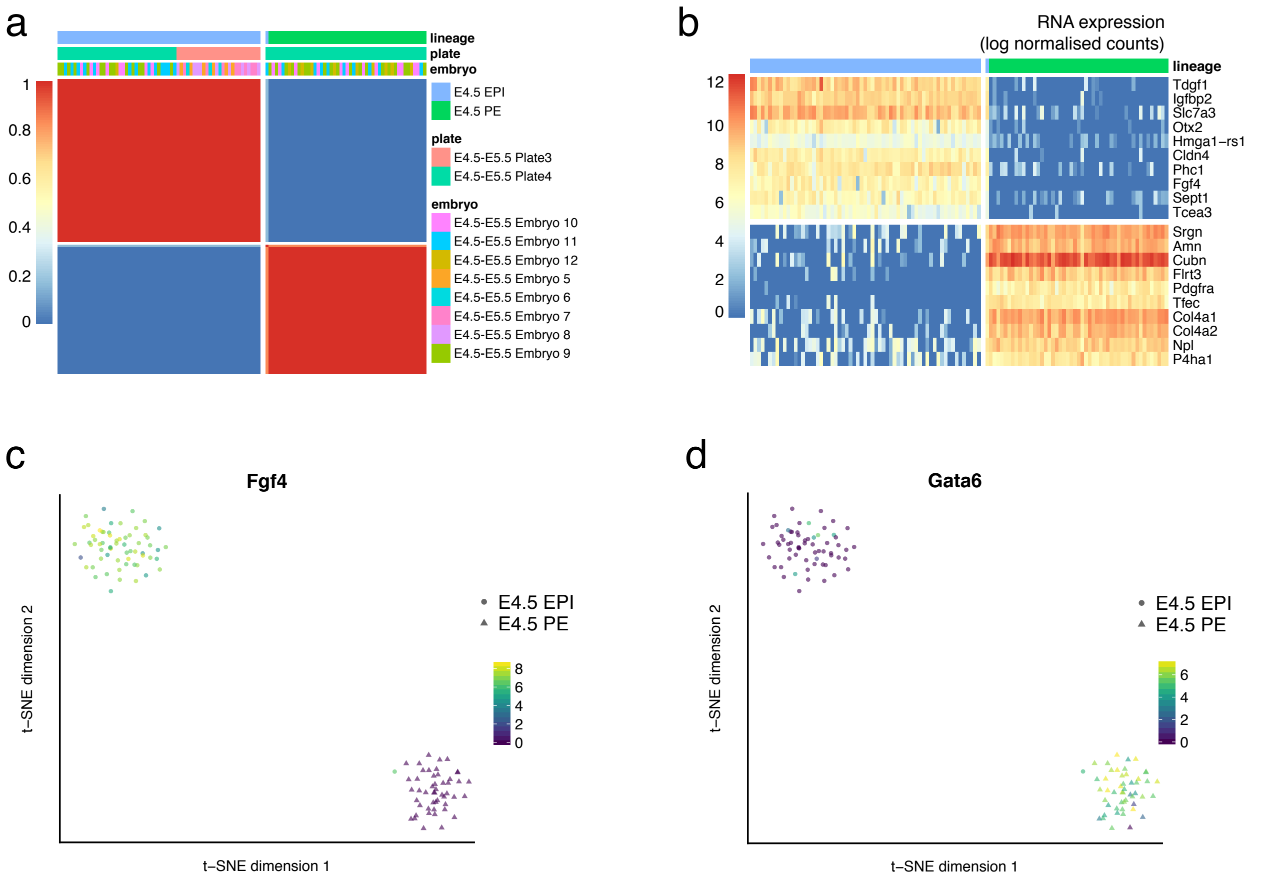


**Figure S2: Characterisation of lineages at E4.5 using SC3.**
(a) Consensus plot for E4.5 representing the similarity between cells based on the averaging of clustering results from all combinations of clustering parameters. Similarity 0 (blue) means that the two cells are always assigned to different clusters, whereas similarity 1 (red) means that the two cells are always assigned to the same cluster. The block-diagonal structure shows the existence of two robust clusters, corresponding to the E4.5 epiblast (E4.5 EPI) and the E4.5 primitive endoderm (E4.5 PE).
(b) Heatmap showing the RNA expression (log normalised counts) of the predicted gene markers for each cluster. For each gene, a binary classifier is constructed based on the mean cluster expression values and an operating characteristic (ROC) curve is used to quantify the accuracy of the prediction. A p-value is assigned to each gene by using the Wilcoxon signed rank test. Top genes were selected using a p-value $<0.01$ and AUROC $>0.85$ thresholds.
(c) t-SNE representation coloured by the RNA expression of *Fgf4*, a known E4.5 epiblast marker[1].
(c) t-SNE representation coloured by the RNA expression of *Gata6*, a known E4.5 primitive endoderm marker[1].

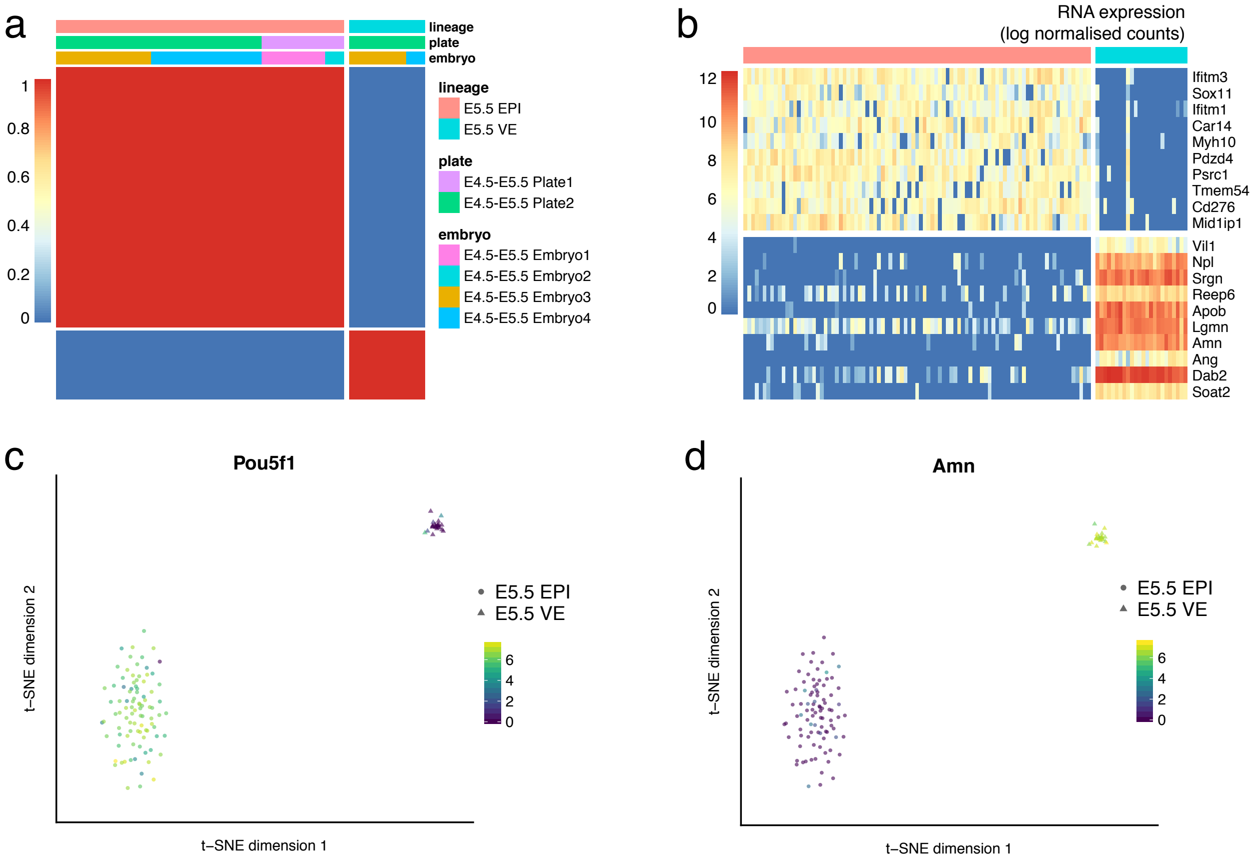


**Figure S3: Characterisation of lineages at E5.5 using SC3.**
(a) Consensus plot for E5.5 representing the similarity between cells based on the averaging of clustering results from all combinations of clustering parameters. Similarity 0 (blue) means that the two cells are always assigned to different clusters, whereas similarity 1 (red) means that the two cells are always assigned to the same cluster. The block-diagonal structure shows the existence of two robust clusters, corresponding to the E5.5 epiblast (E5.5 EPI) and the E5.5 visceral endoderm (E5.5 VE).
(b) Heatmap showing the RNA expression (log normalised counts) of the predicted gene markers for each cluster. For each gene, a binary classifier is constructed based on the mean cluster expression values and an operating characteristic (ROC) curve is used to quantify the accuracy of the prediction. A p-value is assigned to each gene by using the Wilcoxon signed rank test. Top genes were selected using a p-value$<$0.01 and AUROC$>$0.85 thresholds.
(c) t-SNE representation coloured by the RNA expression of *Pou5f1*, a known E5.5 epiblast marker[2].
(d) t-SNE representation coloured by the RNA expression of *Amn*, a known 5.5 visceral endoderm marker[3].

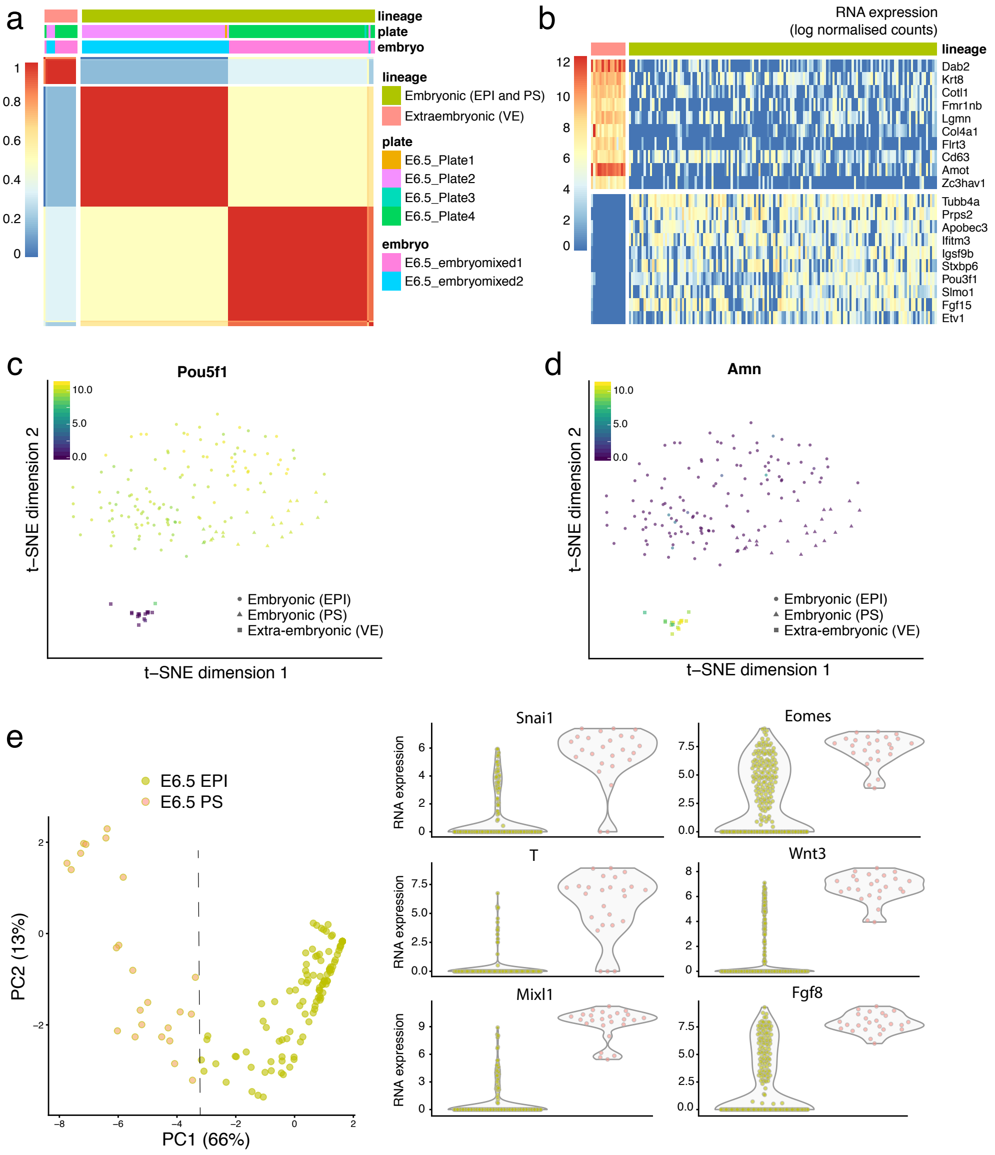


**Figure S4: Characterisation of lineages at E6.5 using SC3 and Principal Component Analysis.**

(a) Consensus plot for E6.5 representing the similarity between cells based on the averaging of clustering results from all combinations of clustering parameters. Similarity 0 (blue) means that the two cells are always assigned to different clusters, whereas similarity 1 (red) means that the two cells are always assigned to the same cluster. The block-diagonal structure shows the existence of at least two robust clusters, corresponding to embryonic cells (E6.5 epiblast (E6.5 EPI) and E6.5 primitive streak (E6.5 PS), and non-embryonic cells (E6.5 visceral endoderm (E6.5 VE)).

(b) Heatmap showing the RNA expression (log normalised counts) of the predicted gene markers for each cluster. For each gene, a binary classifier is constructed based on the mean cluster expression values and an operating characteristic (ROC) curve is used to quantify the accuracy of the prediction. A p-value is assigned to each gene by using the Wilcoxon signed rank test. Top genes were selected using a p-value$<$0.01 and AUROC$>$0.85 thresholds.

(c) t-SNE representation coloured by the RNA expression for *Pou5f1*, a known epiblast marker[2].

(d) t-SNE representation coloured by the RNA expression for *Amn*, a known visceral endoderm marker[3].

(e) Unsupervised SC3 clustering was not able to identify primitive streak cells. Instead, we used Principal Component Analysis on a list of selected marker genes: *Snai1*, *T*, *Mixl*1, *Eomes*, *Wnt3*, *Fgf8*[4]. Left, scatter plot of the two first principal components of the subsetted RNA expression matrix. A threshold on PC1 (dashed line) was used to classify E6.5 epiblast from E6.5 primitive streak cells. Right, RNA expression (log counts) of the selected marker genes.

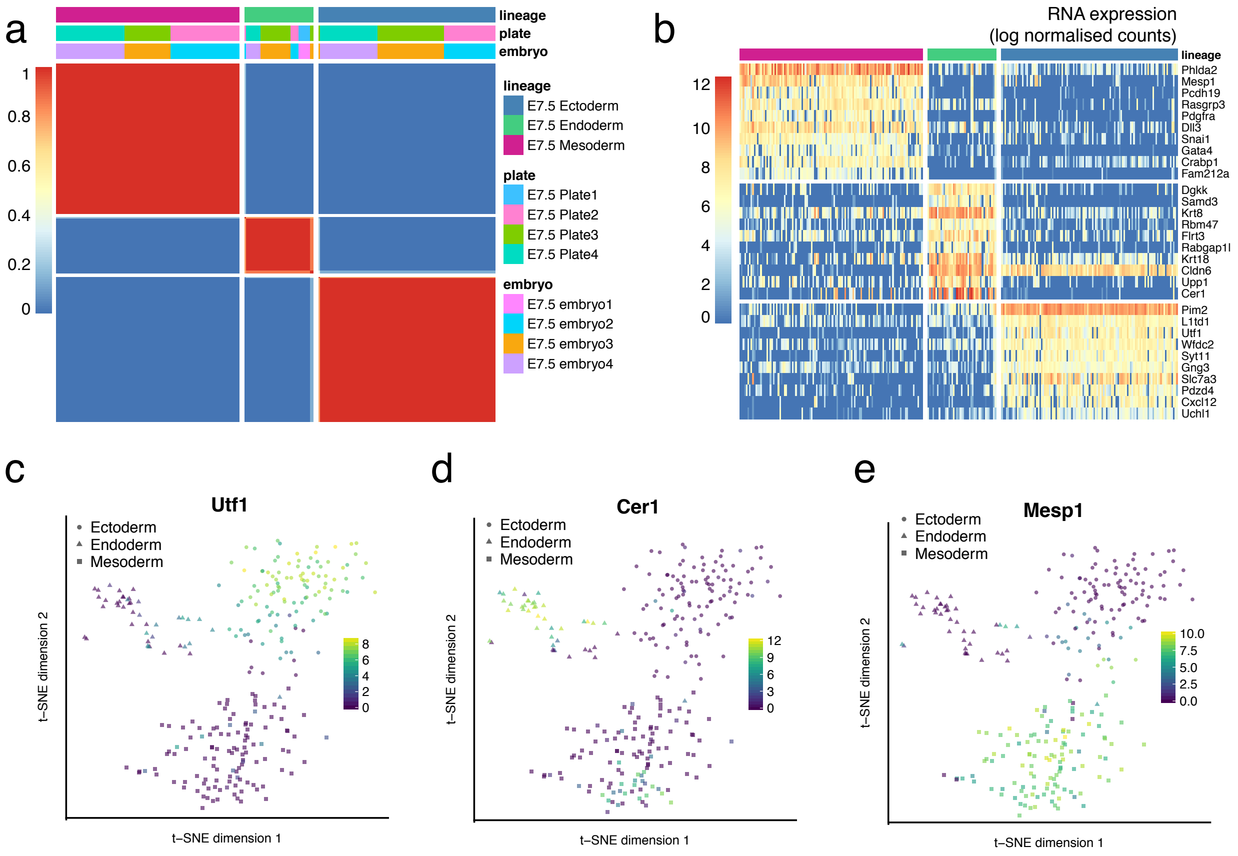


**Figure S5: Characterisation of lineages at E7.5 using SC3.**
(a) Consensus plot for E7.5 representing the similarity between cells based on the averaging of clustering results from all combinations of clustering parameters. Similarity 0 (blue) means that the two cells are always assigned to different clusters, whereas similarity 1 (red) means that the two cells are always assigned to the same cluster. The block-diagonal structure shows the existence of three robust clusters, corresponding to the three germ layers: E7.5 ectoderm, E7.5 mesoderm and E7.5 endoderm.
(b) Heatmap showing the RNA expression (log normalised counts) of the predicted gene markers for each cluster. For each gene, a binary classifier is constructed based on the mean cluster expression values and an operating characteristic (ROC) curve is used to quantify the accuracy of the prediction. A p-value is assigned to each gene by using the Wilcoxon signed rank test. Top genes were selected using a p-value $<0.01$ and AUROC $>0.85$ thresholds.
(c) t-SNE representation coloured by the RNA expression of *Utf1*, a known ectoderm marker[5].
(d) t-SNE representation coloured by the RNA expression of *Cer1*, a known endoderm marker[6].
(e) t-SNE representation coloured by the RNA expression of *Mesp1*, a known mesoderm marker[7].

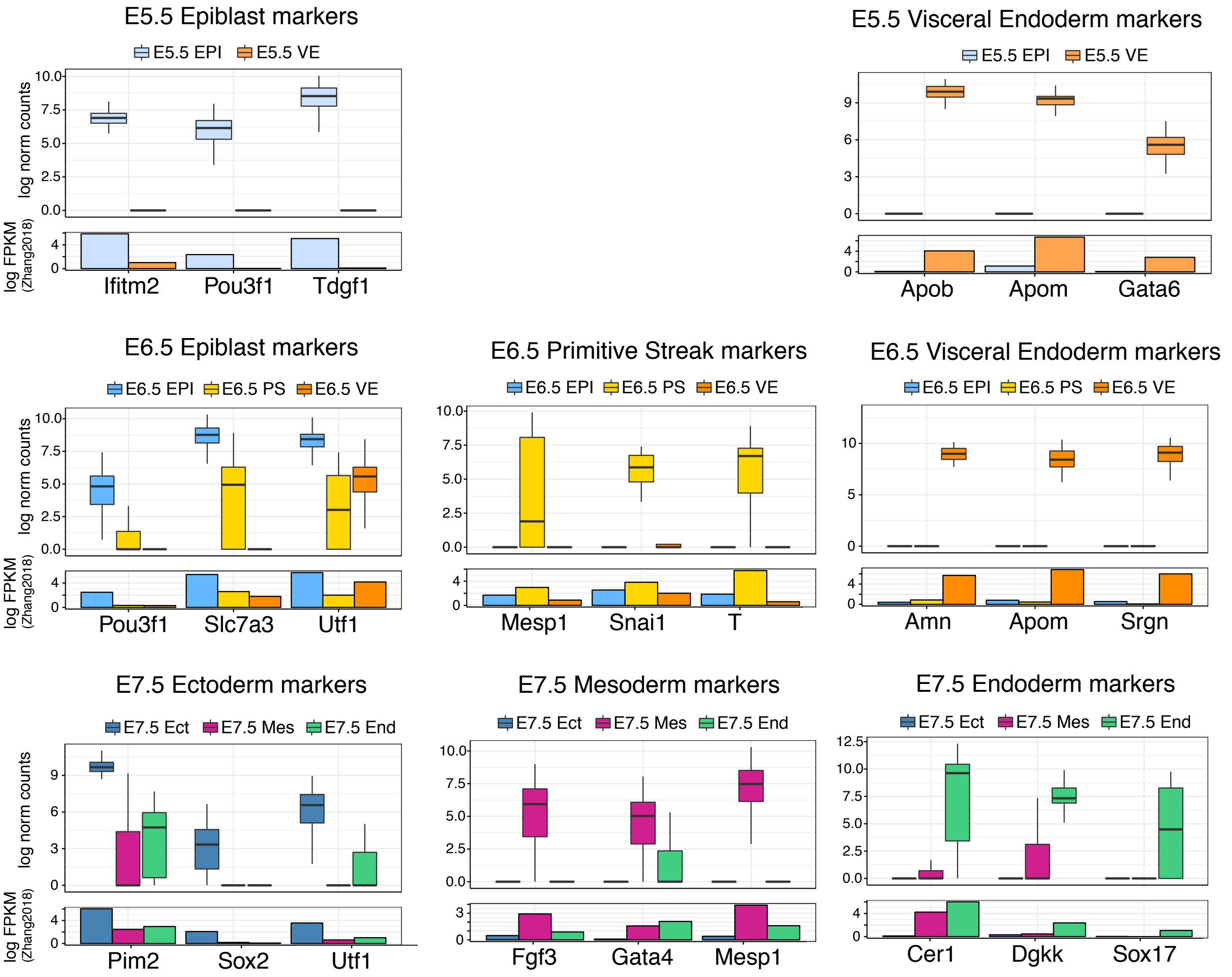


**Figure S6: Validation of lineage-specific RNA expression markers.**
Shown is the gene expression of selected markers for each stage and lineage. The box plots represent the distribution of log2 normalised counts from this study (each data point is a single cell) and the bar plots show the bulk log FPKM values from[8].

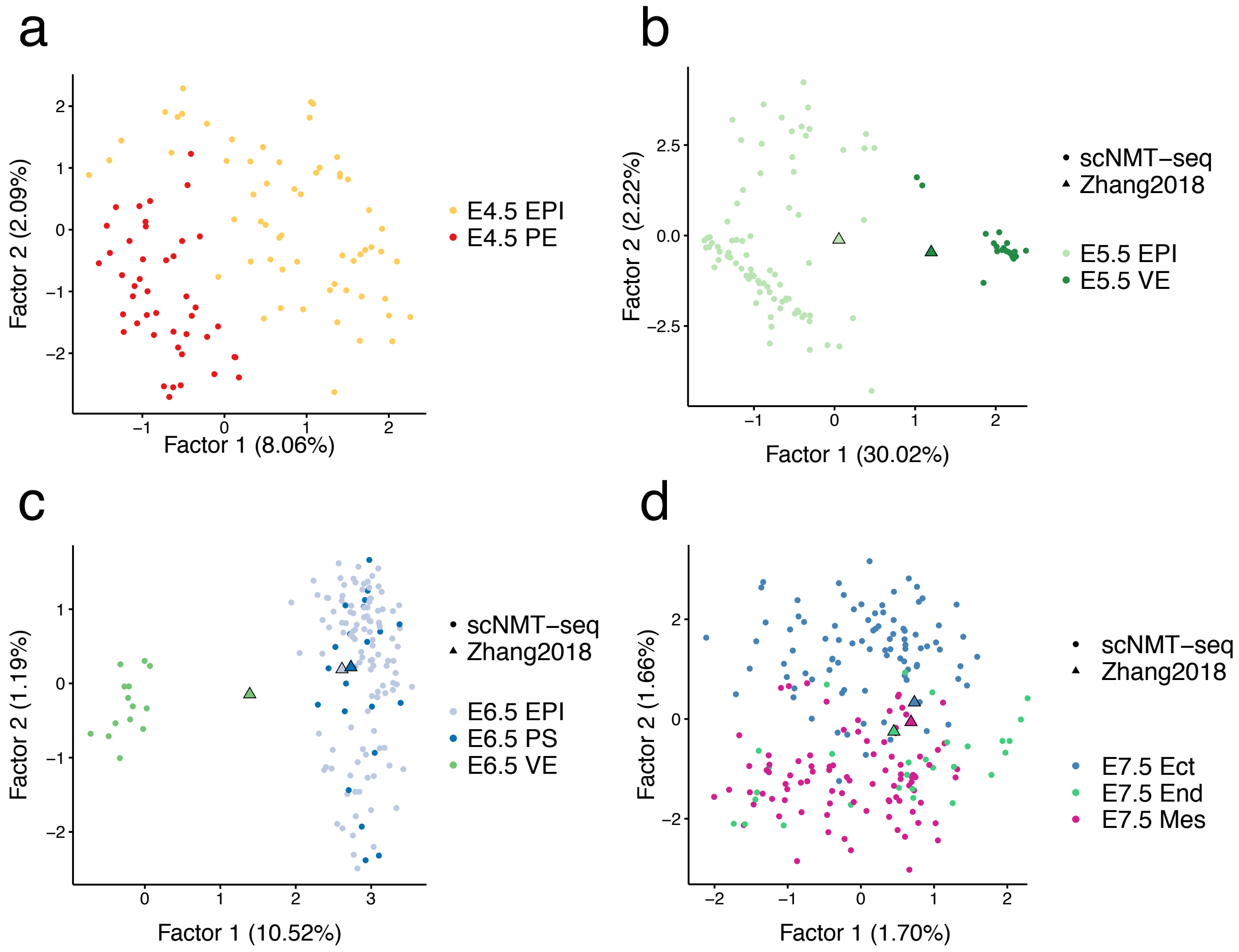


**Figure S7: Unsupervised dimensionality reduction of DNA methylation data separates embryonic from extraembryonic cells.**
To perform dimensionality reduction while handling the large amount of missing values we used a Bayesian Factor Analysis model[9]. One model was trained per stage. Shown are scatter plots of the first two latent factors (sorted by variance explained) for (a) E4.5, (b) E5.5, (c) E6.5 and (d) E7.5, coloured by lineage (assigned based on the RNA expression profiles). The fraction of variance explained by each factor is displayed in parentheses. Where available, we included bulk data from[8] (triangles).
The input data was DNA methylation levels (%) quantified using a running window of 5kb across the genome. To increase computational efficiency we selected the top 5,000 most variable sites across cells for training the model.

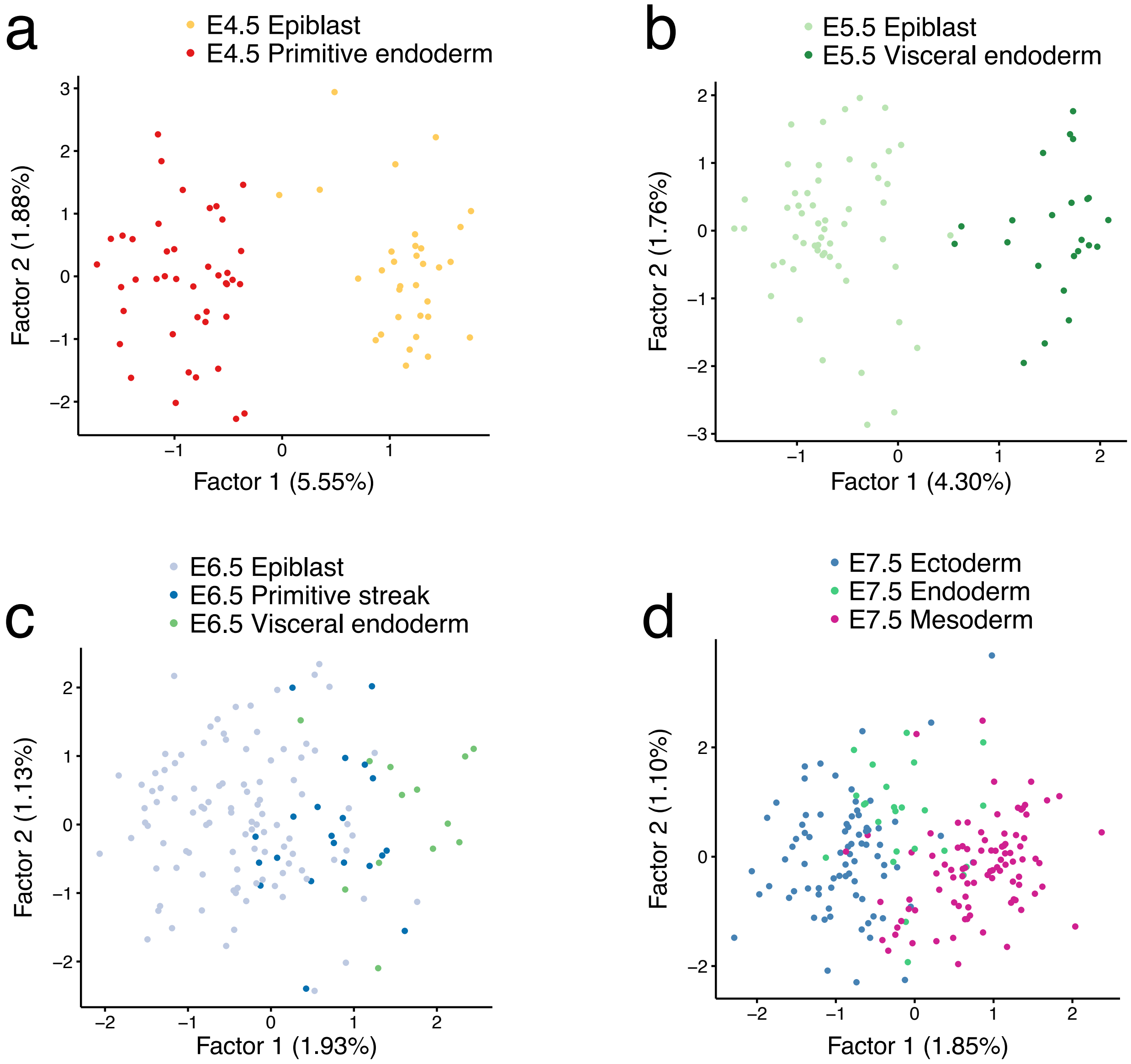


**Figure S8: Unsupervised dimensionality reduction of chromatin accessibility data separates embryonic from extraembryonic cells.**
To perform dimensionality reduction while handling the large amount of missing values we used a Bayesian Factor Analysis model[9]. One model was trained per stage. Shown are scatter plots of the first two latent factors (sorted by variance explained) for the different stages: (a) E4.5, (b) E5.5, (c) E6.5 and (d) E7.5. In parentheses, we show the fraction of variance explained per factor.
The input data was chromatin accessibility rates quantified in non-overlapping 100bp windows. To select informative sites and increase computational efficiency, we first selected accessible (>75%) 100bp windows in pseudobulk E4.5 data (≈250,000 sites) then filtered for the most variable sites (top 5,000) at a given stage.

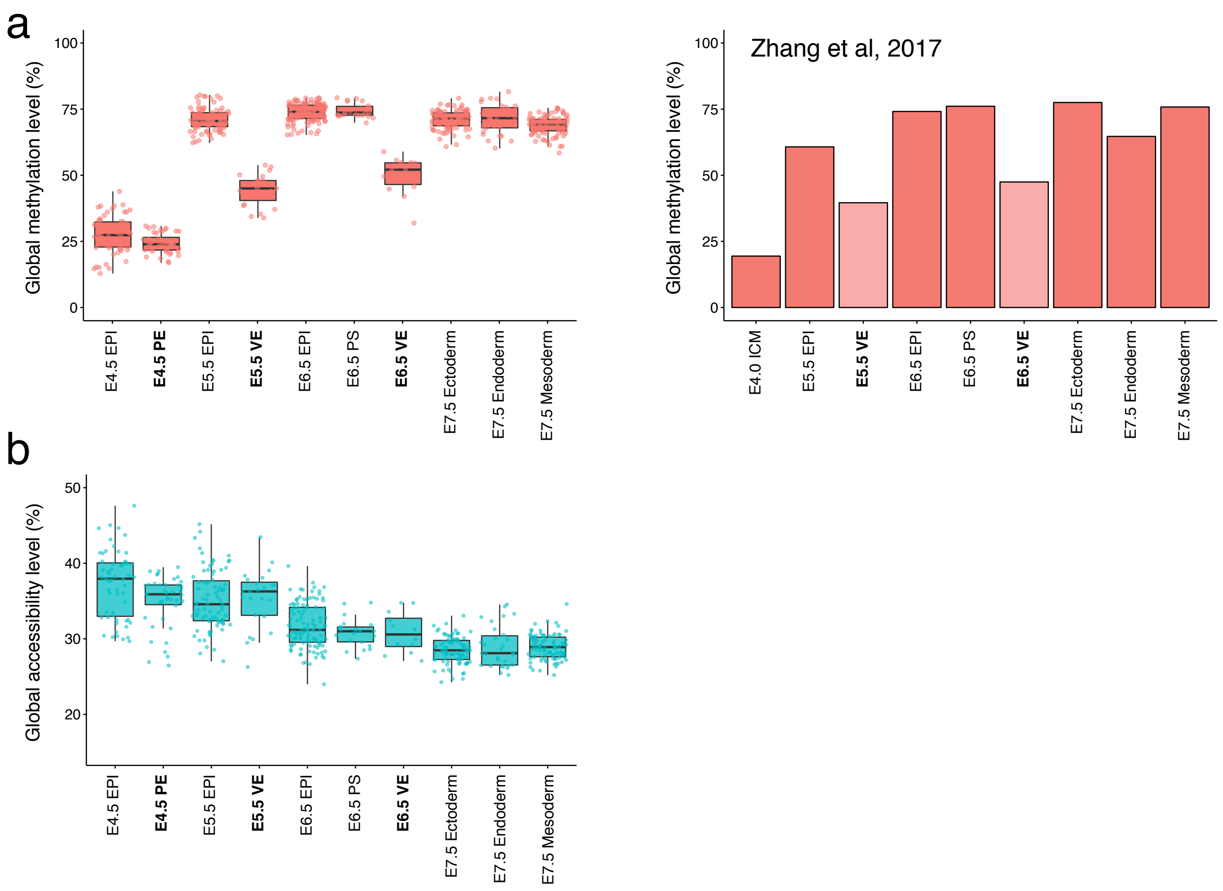


**Figure S9: Global DNA methylation and chromatin accessibility statistics per lineage.**
(a) Left: boxplots showing the distribution of genome-wide CpG methylation levels (%) per stage and lineage. Each dot depicts a single cell from this study. Right: barplots showing the genome-wide CpG methylation levels (%) per stage and lineage from published bulk data[8].
(b) Boxplots showing the distribution of genome-wide GpC accessibility levels (%) per stage and lineage. Each dot depicts a single cell from this study.

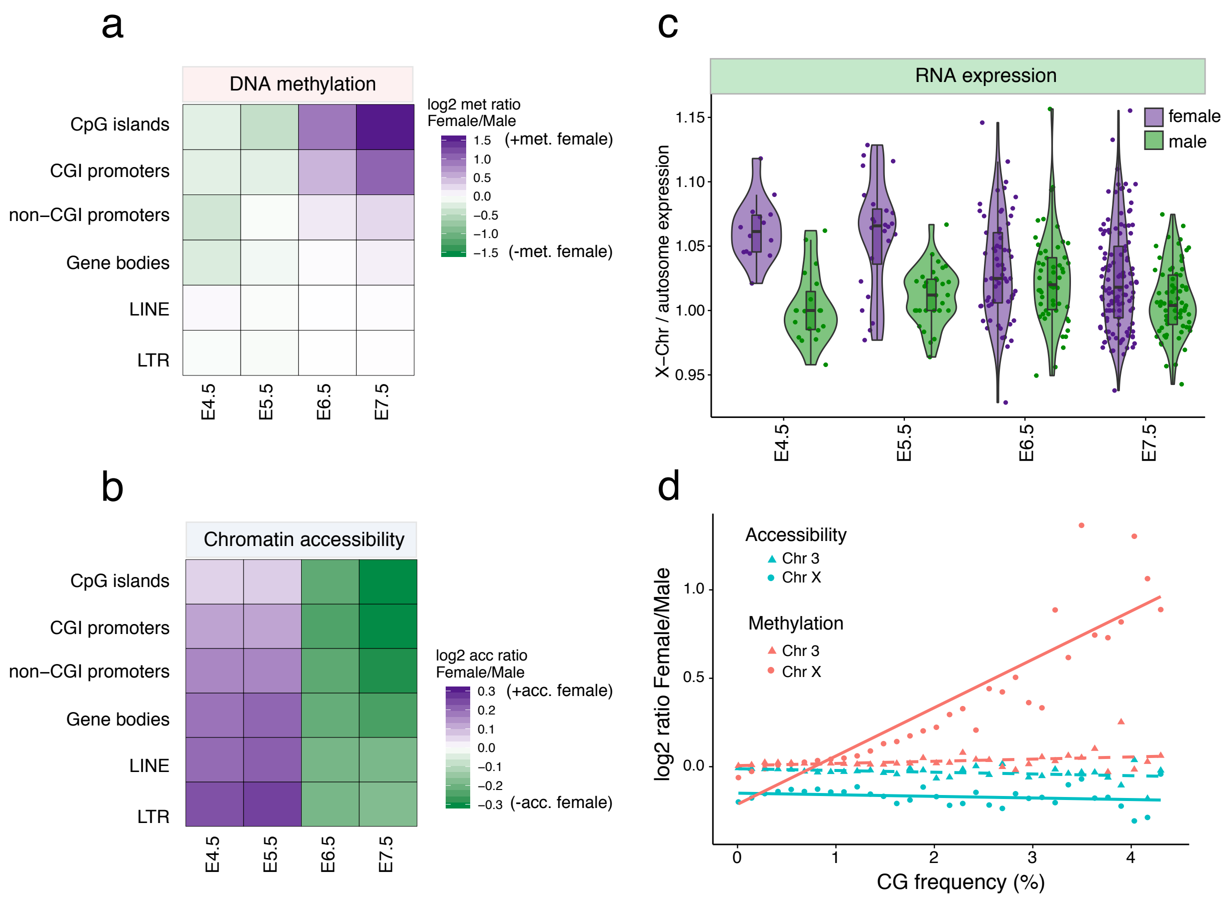


**Figure S10: RNA expression, chromatin accessibility and DNA methylation dynamics associated with X chromosome inactivation.**
(a) Heatmap of the mean log2 female to male DNA methylation ratio on the X chromosome at different genomic contexts, sorted by CpG density. Purple squares represent higher methylation levels in female cells, green squares represent higher methylation levels in male cells.
(b) Heatmap of the mean log2 female to male chromatin accessibility ratio on the X chromosome at different genomic contexts, sorted by CpG density. Purple squares represent higher accessibility levels in female cells, green squares represent higher accessibility levels in male cells.
(c) Median X chromosome RNA expression level normalised over median autosomal RNA expression level, for female (purple) and male (green) cells. Each dot represents a single cell. At E4.5 and E5.5 X chromosome expression is higher in female cells, indicating that both female copies of the X chromosome are actively transcribed. From E6.5 X chromosome expression is similar in female and male cells indicating that one copy of the female X chromosome is silenced.
(d) Scatter plot showing the relationship between CpG density and female to male log2 DNA methylation (red) or chromatin accessibility (blue) ratio for the X chromosome (circles) and chromosome 3 (triangles, control). Each chromosome was divided in 2kb bins, and CpG density, mean female methylation/accessibility rate and mean male methylation/accessibility rate were calculated for each bin. For plotting purposes, bins were further aggregated by CpG density (100 bins).

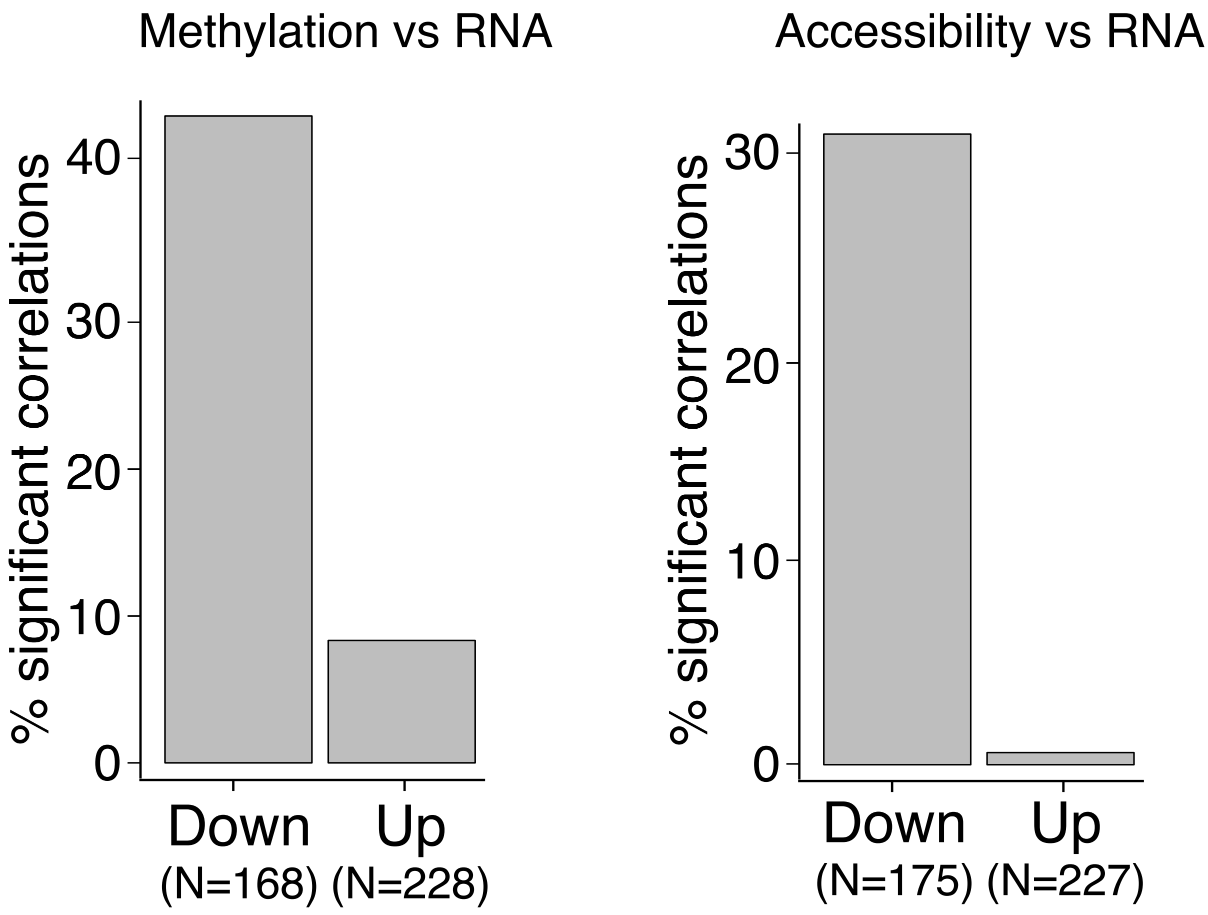


**Figure S11: Epigenetic changes in promoters are associated with the repression of early pluripotency markers.**

Bar plots show the fraction of genes with significant association between DNA methylation versus RNA expression (left), and chromatin accessibility versus RNA expression (right). Genes are split according to whether they are up-regulated (Up, greater expression in E7.5) or down-regulated (Down, greater expression in E4.5).

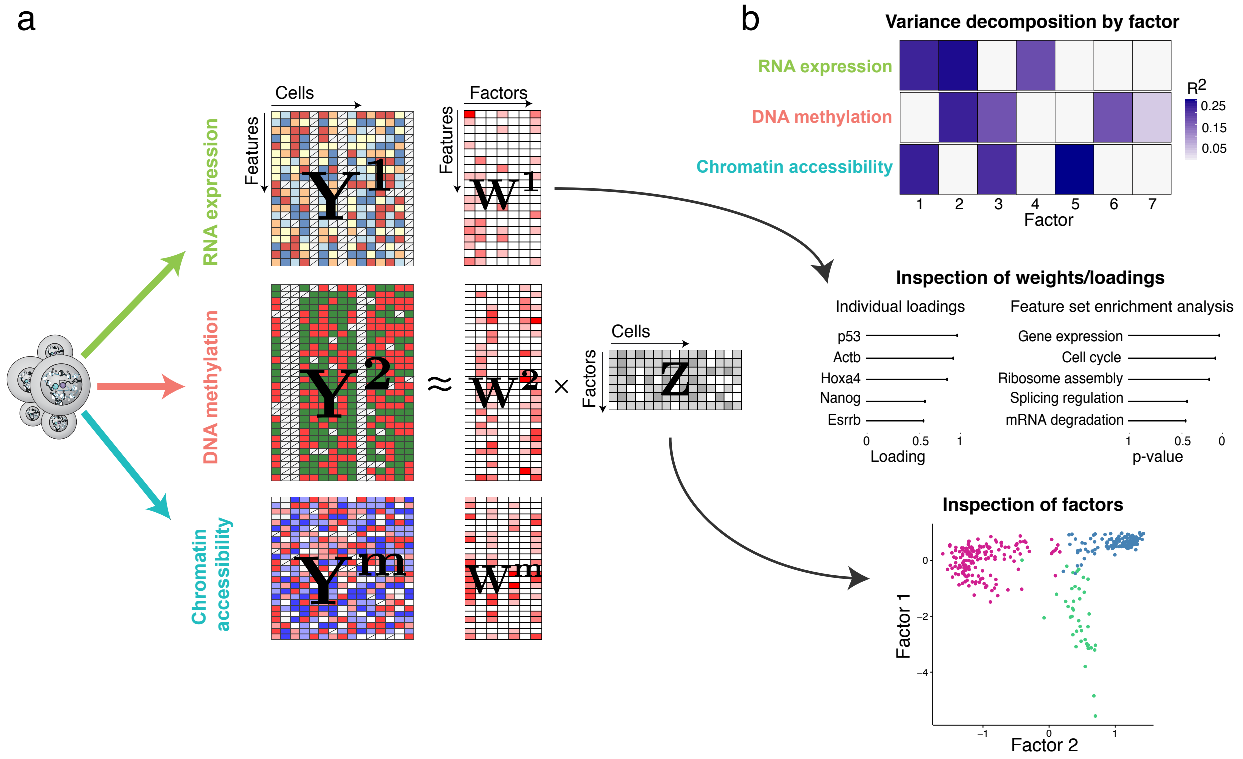


**Figure S12: Multi-Omics Factor Analysis (MOFA): model overview and illustration of downstream analysis.**
(a) Model overview: MOFA takes as input one or more data matrices (Y) from each data modality, extracted from the same single cells. MOFA decomposes these matrices into a matrix of factors (Z) and M weight matrices (W), one for each data modality. The Z matrix contains the low dimensional representation of cells in terms of latent factors. The W matrices relate the low-dimensional space to the high-dimensional space by inferring a loading for each feature on each factor. When interpreting a factor, the absolute value of the loading is used as a measure of feature importance.
(b) Downstream analysis: the fitted MOFA model can be queried for different downstream analyses, including (i) variance decomposition, assessing the proportion of variance ($R^2$) explained by each factor in each data modality, (ii) semi-automated factor annotation based on the inspection of loadings and gene set enrichment analysis, (iii) visualization of the samples in the factor space.

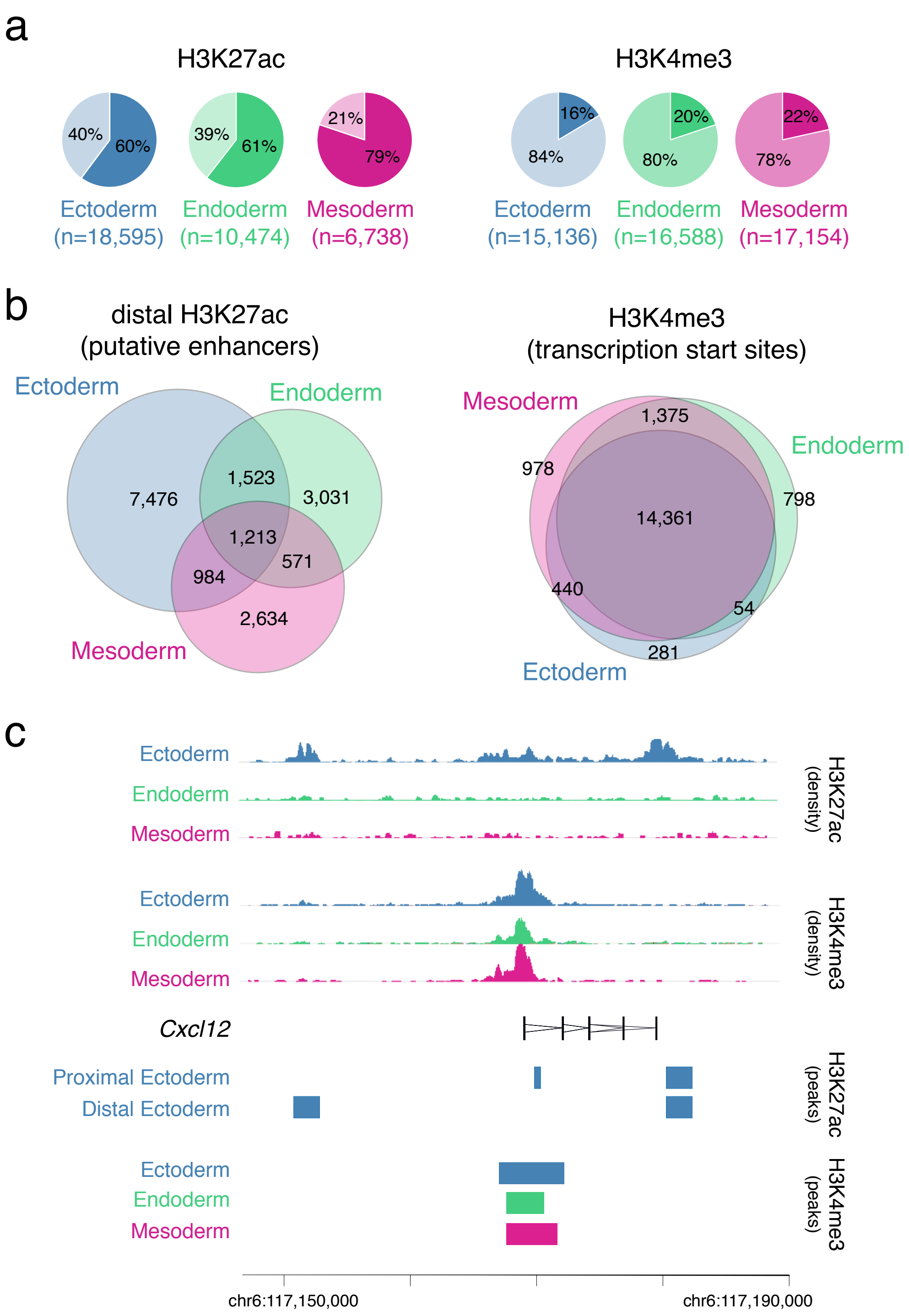


**Figure S13: Characterisation of lineage-specific H3K27ac and H3K4me3 ChIP-seq data.** Peaks were called individually for each lineage and histone mark (see Methods).

(a) Percentage of peaks overlapping promoters (+/- 500 bp of transcriptional start sites of annotated Ensembl mRNAs; lighter colour) and not overlapping promoters (distal peaks, darker colour).

(b) Venn diagrams showing overlap of peaks for each lineage, for distal H3K27ac (left) and all H3K4me3 (right). This shows that H3K27ac peaks tend to be distal from the promoters and those tend to be lineage-specific, marking putative enhancer elements[10]. In contrast, H3K4me3 peaks tend to be shared between lineages and overlap promoter regions, marking transcription start sites[11].

(c) Illustrative example of the ChIP-seq profile of the *Cxcl12* gene. The top tracks show wiggle plots of ChIP-seq read density (normalised by total read count) for lineage-specific H3K27ac and H3K4me3. The coding sequence is shown in black. The bottom tracks show the lineage-specific peak calls (see Methods). H3K27ac are split into distal (putative enhancers) and proximal to the promoter.

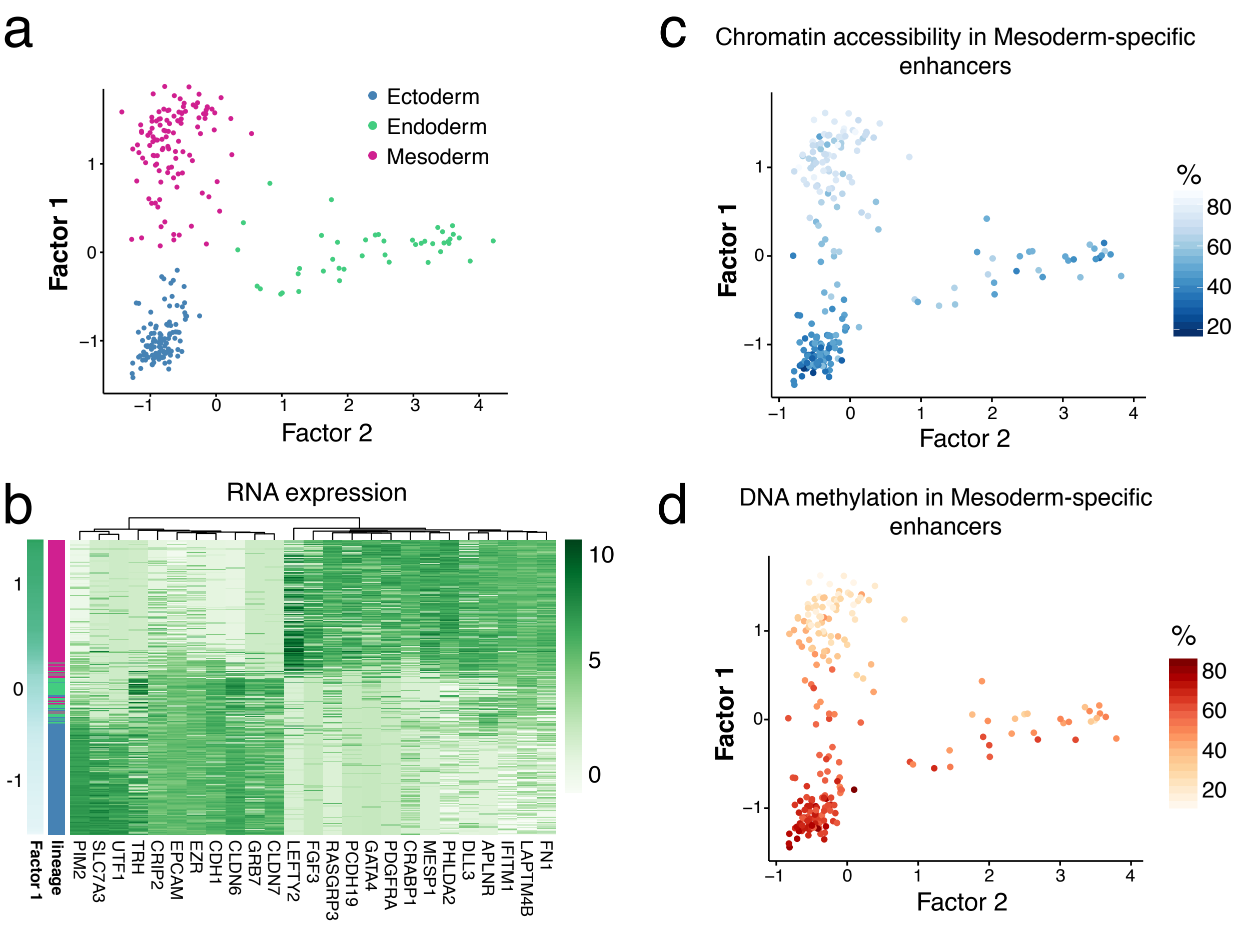


**Figure S14: Characterisation of MOFA Factor 1 as the mesoderm formation factor.**
(a) Scatter plot of Factor 2 (x-axis) and Factor 1 (y-axis) values, coloured by lineage.
(b) Heatmap of the RNA expression profiles for the top genes with highest loading. Genes are grouped by hierarchical clustering. Cells are sorted according to Factor 1 values, from more mesoderm-like (top) to less mesoderm-like (bottom).
(c-d) Scatter plot of Factor 2 (x-axis) and Factor 1 (y-axis) values, coloured by (d) mean chromatin accessibility and (e) mean DNA methylation levels (%) of the top 100 mesoderm-specific enhancers with highest loading.

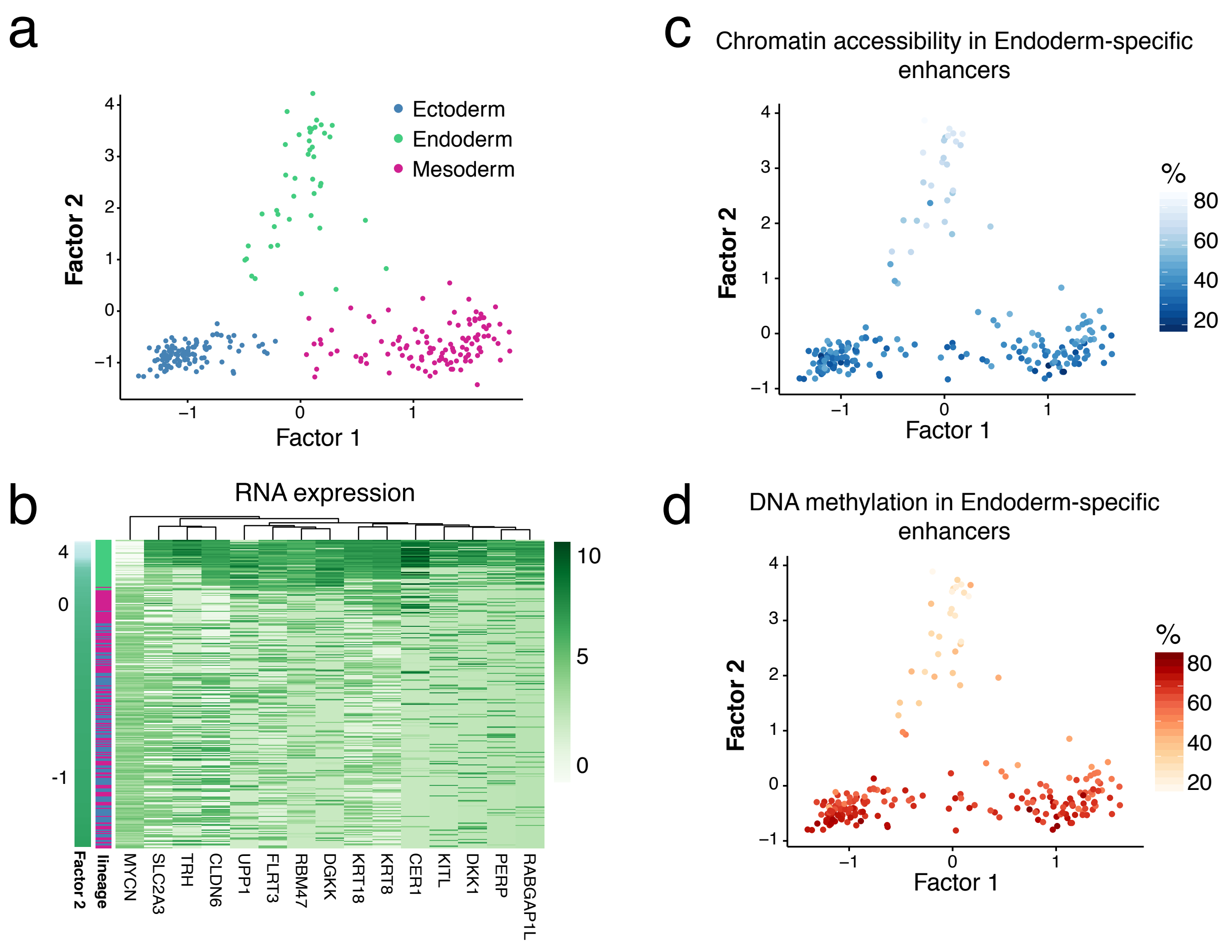


**Figure S15: Characterisation of MOFA Factor 2 as the endoderm formation factor.**
(a) Scatter plot of Factor 1 (x-axis) and Factor 2 (y-axis) values, coloured by lineage.
(b) Heatmap of the RNA expression profiles for the genes with highest loading. Genes are grouped by hierarchical clustering. Cells are sorted according to Factor 2 values, from more endoderm-like (top) to less endoderm-like (bottom).
(c-d) Scatter plot of Factor 1 (x-axis) and Factor 2 (y-axis) values, coloured by (d) mean chromatin accessibility and (e) mean DNA methylation levels (%) of the top 100 endoderm-specific enhancers with highest loading.

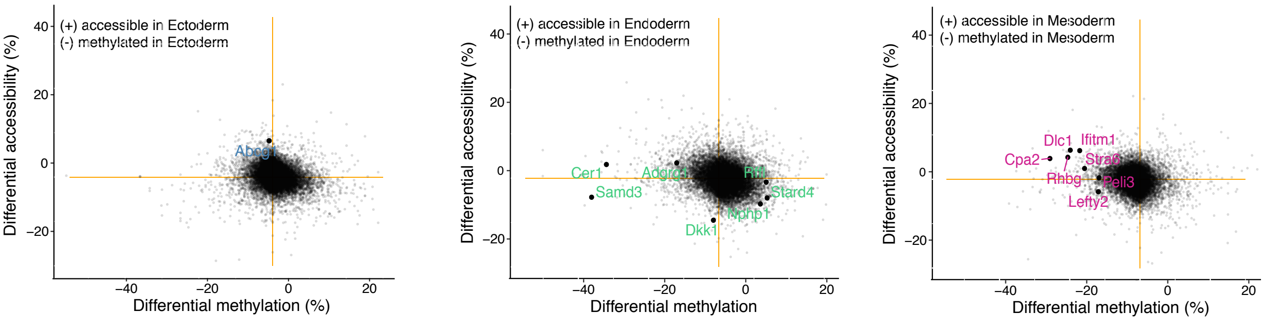


**Figure S16: Differential DNA methylation and chromatin accessibility analysis in promoters at E7.5.**

Scatter plots showing differential DNA methylation (x-axis) versus chromatin accessibility (y-axis) analysis at promoters. Each dot corresponds to a gene. Shown are ectoderm vs non-ectoderm cells (left), endoderm vs non-endoderm cells (middle) and mesoderm vs non-mesoderm cells (right). Labeled big dots depict genes with lineage-specific RNA expression that show significant differential methylation or accessibility in their promoter (FDR<10%).

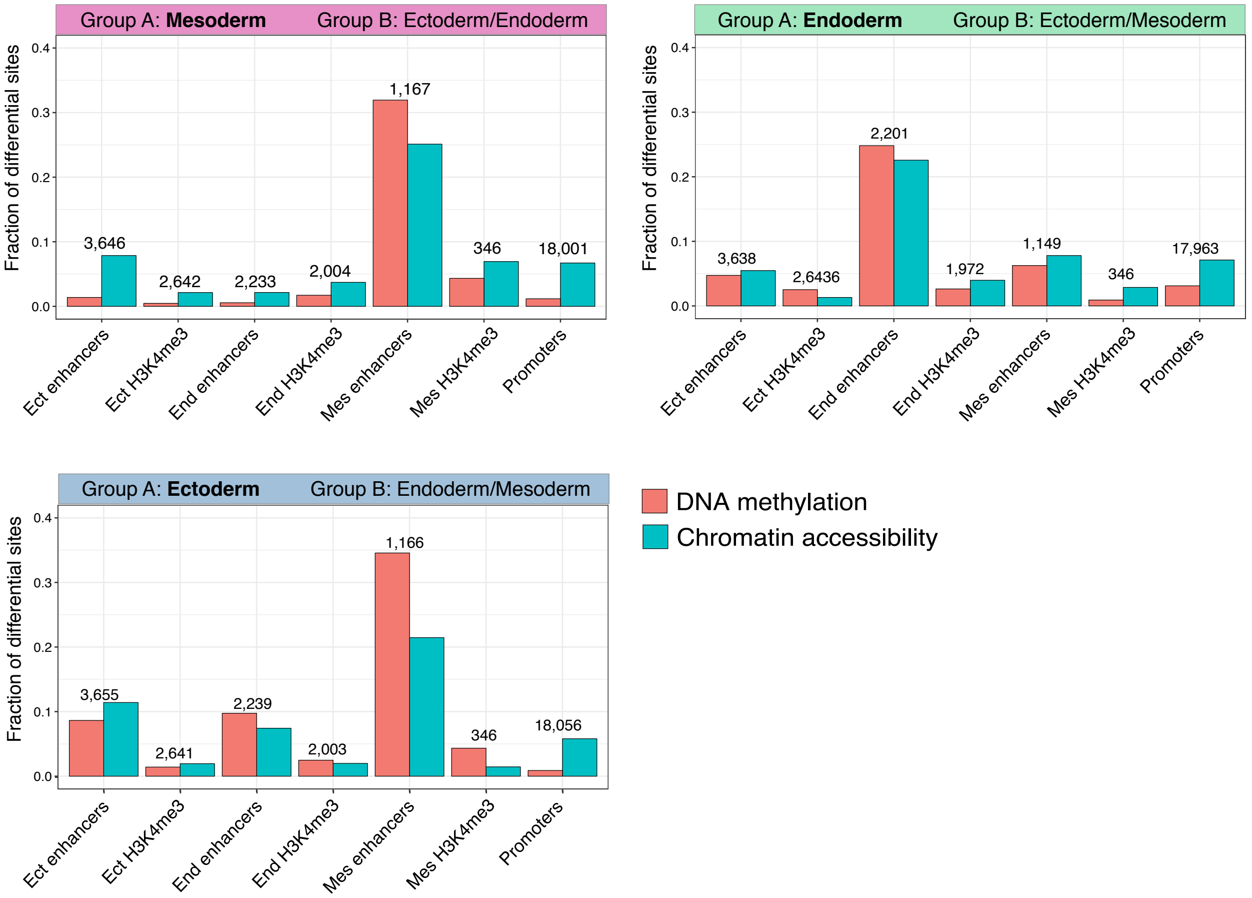


**Figure S17: Differential DNA methylation and chromatin accessibility analysis at E7.5 for different genomic contexts.**

Bar plots show the fraction of differentially methylated (red) or accessible (blue) loci (FDR<10%, y-axis) per genomic context (x-axis). Each subplot corresponds to the comparison of one lineage against cells comprising the other lineages present at E7.5.

Differential analysis of DNA methylation and chromatin accessibility was performed independently for each genomic element using a Fisher exact test of equal proportions (see Methods). The number on top of each bar plot shows the number of loci that were tested.

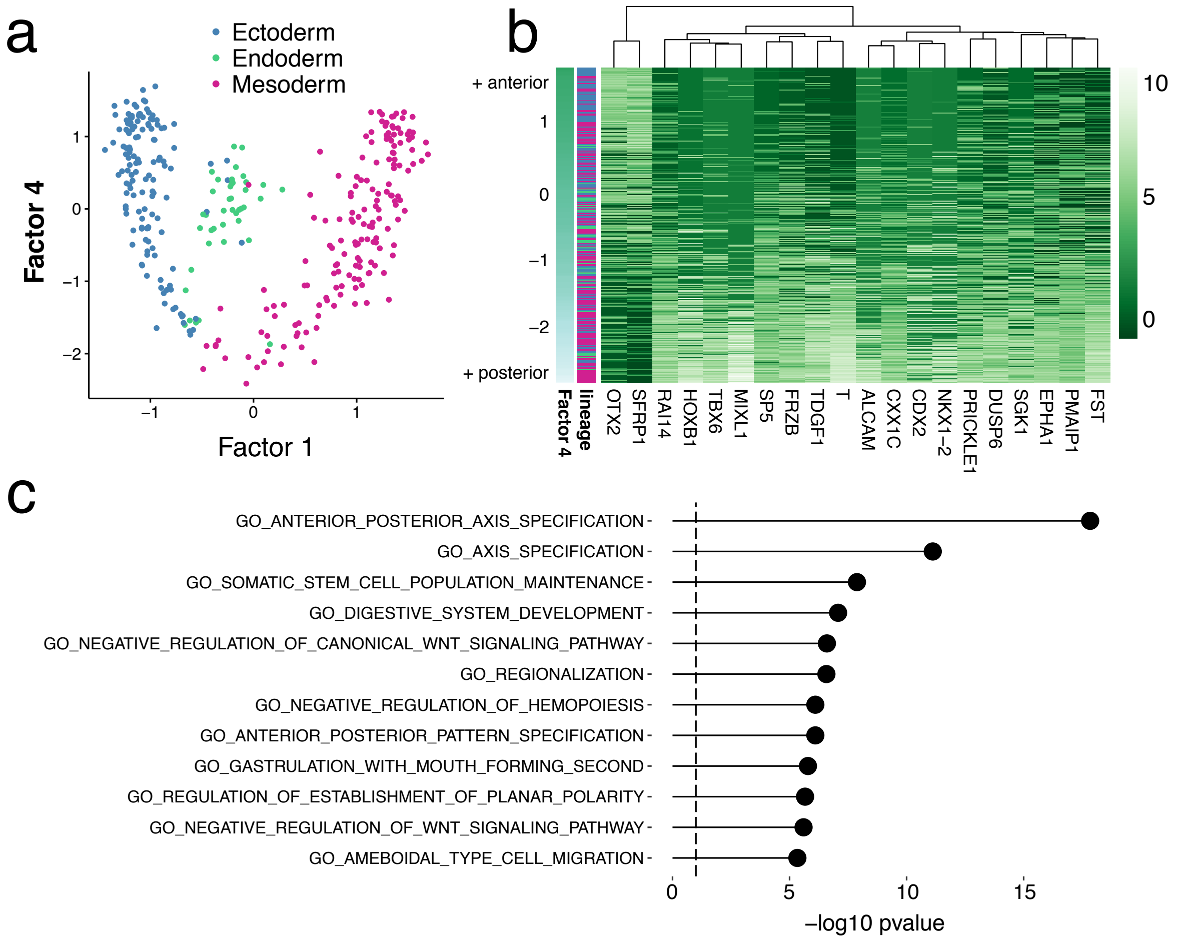


**Figure S18: Characterisation of MOFA Factor 4 as antero-posterior axial patterning.**
(a) Scatter plot of Factor 1 (x-axis) and Factor 4 (y-axis) values, coloured by lineage.
(b) Heatmap displaying the RNA expression profiles for the genes with highest loading on Factor 4. Genes are grouped by hierarchical clustering. Samples are sorted according to the factor values from more anterior (top) to more posterior (bottom).
(c) Gene set enrichment analysis of the gene loadings of Factor 4. Shown are the top most significant pathways from MSigDB C2[12,13].

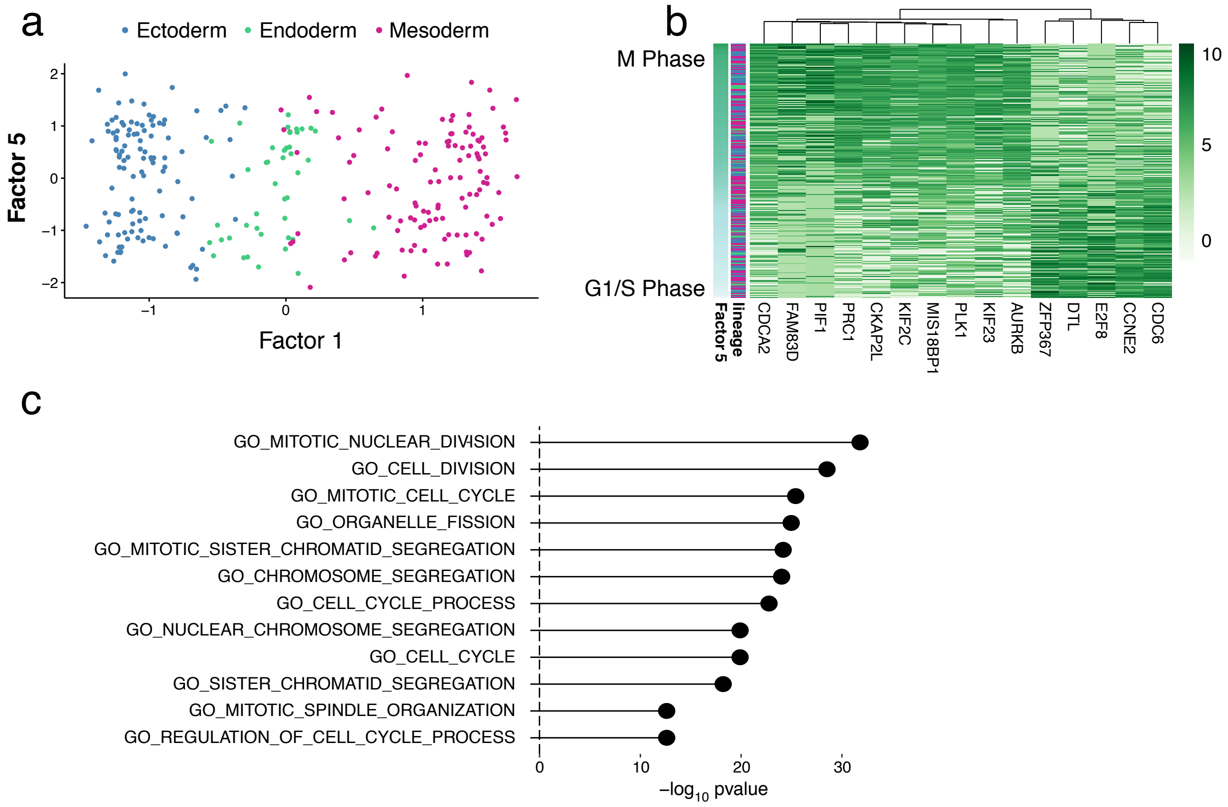


**Figure S19: Characterisation of MOFA Factor 5 as cell cycle.**
(a) Scatter plot of Factor 1 (x-axis) and Factor 5 (y-axis) values, coloured by lineage.
(b) Heatmap displaying the RNA expression profiles for the genes with highest loading on Factor 5. Genes are grouped by hierarchical clustering. Samples are sorted according to the factor values from M phase (top) to G1/S phase(bottom).
(c) Gene set enrichment analysis of the gene loadings of Factor 5. Shown are the top most significant pathways from MSigDB C2[12,13].

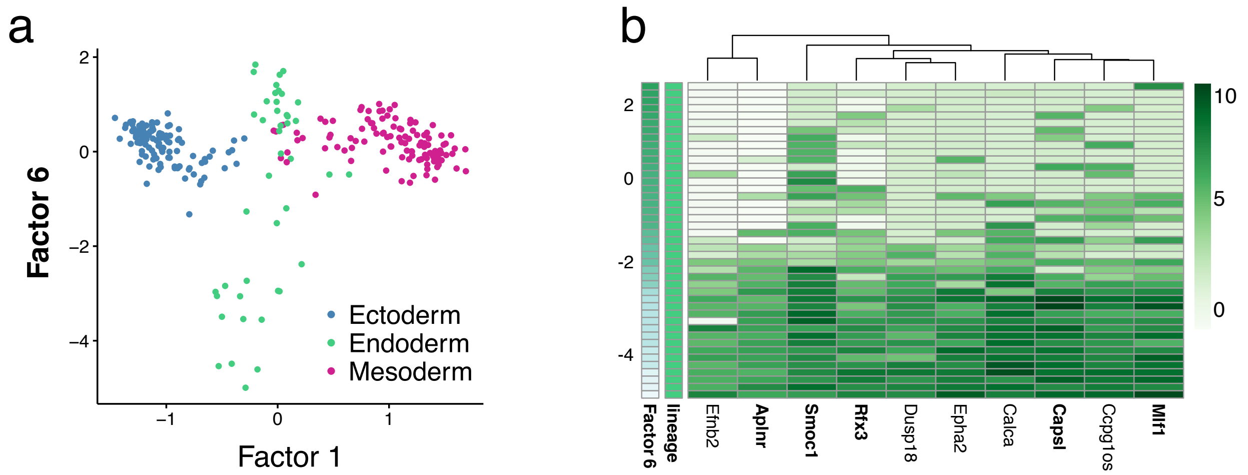


**Figure S20: Characterisation of MOFA Factor 6 as node proximity and notochord formation in the endoderm.**
(a) Scatter plot of Factor 1 (x-axis) and Factor 6 (y-axis) values, coloured by lineage.
(b) Heatmap displaying the RNA expression profiles for the genes with highest loading on Factor 6. Genes are grouped by hierarchical clustering. Samples are sorted according to the factor values from less notochord-like (top) to more notochord-like (bottom). In bold, genes with a known association to notochord formation and/or node proximity[14–16].

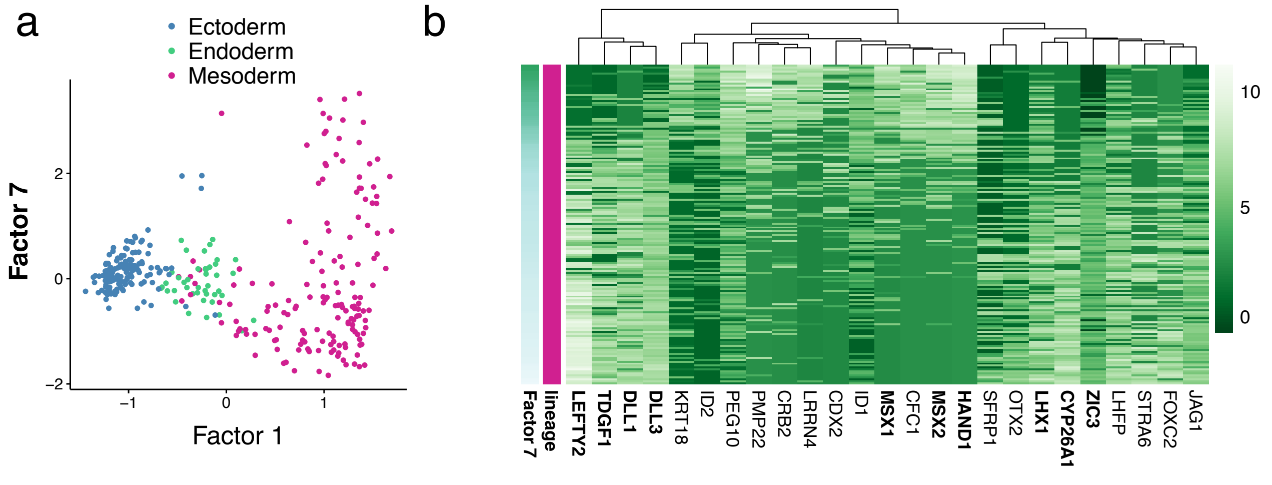


**Figure S21: Characterisation of MOFA Factor 7 as mesoderm patterning.**
(a) Scatter plot of Factor 1 and Factor 7 values, coloured by lineage.
(b) Heatmap displaying the RNA expression profiles for the genes with highest loading on Factor 7. Genes are grouped by hierarchical clustering. Samples are sorted according to the factor values. Bolded are genes involved in mesoderm patterning and maturation, according to the MGI Gene Expression database[17].

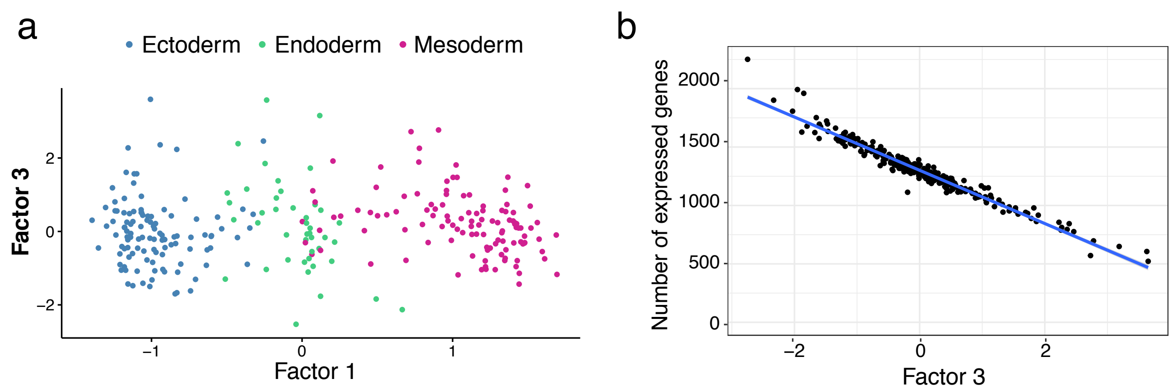


**Figure S22: Characterisation of MOFA Factor 3 as the cellular detection rate.**
(a) Scatter plot of Factor 1 (x-axis) and Factor 3 (y-axis) values, coloured by lineage.
(b) Scatter plot of Factor 3 values (x-axis) and number of expressed genes per cell (y-axis). The blue line shows the linear regression fit.

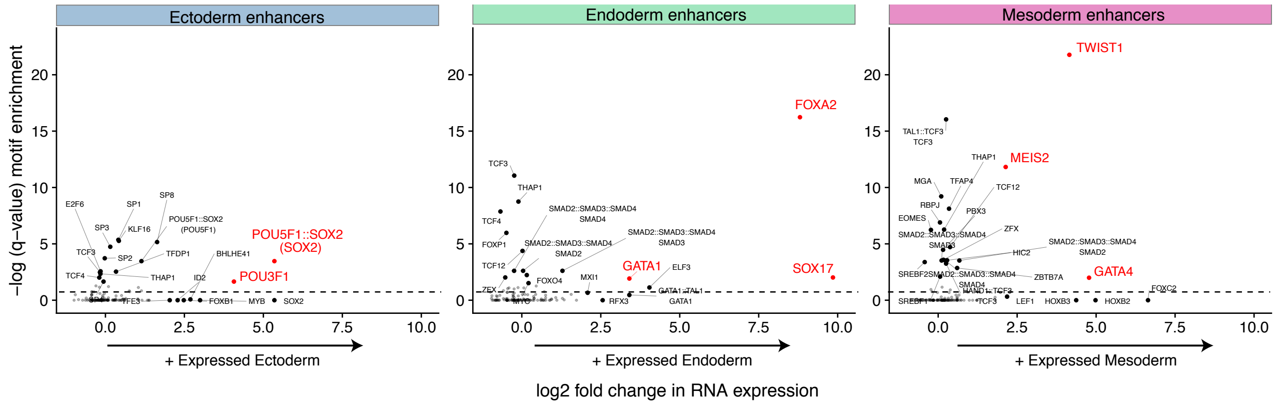


**Figure S23: Identification of transcription factors enriched in lineage-specific enhancers during germ layer commitment.**
To find TF motifs enriched in lineage-associated sites, we used lineage-specific H3K27ac sites that were identified as differentially accessible (in each corresponding lineage) and tested for enrichment over a background of all H3K27ac sites (see Methods).
Shown is motif enrichment (-log10 FDR corrected p-value, y-axis) plotted against differential RNA expression (log fold change, x-axis) of the corresponding transcription factor (TF). The analysis is performed separately for each germ-layer.
TFs significant for either enrichment (FDR<5%) or expression (FDR<5% and log-fold change higher than 2) are coloured in black and labelled. Those significant for both are coloured in red. When a motif is associated with multiple TFs, including heterodimers, we plot the motifs separately and label in brackets the TF that is used for testing differential RNA expression.

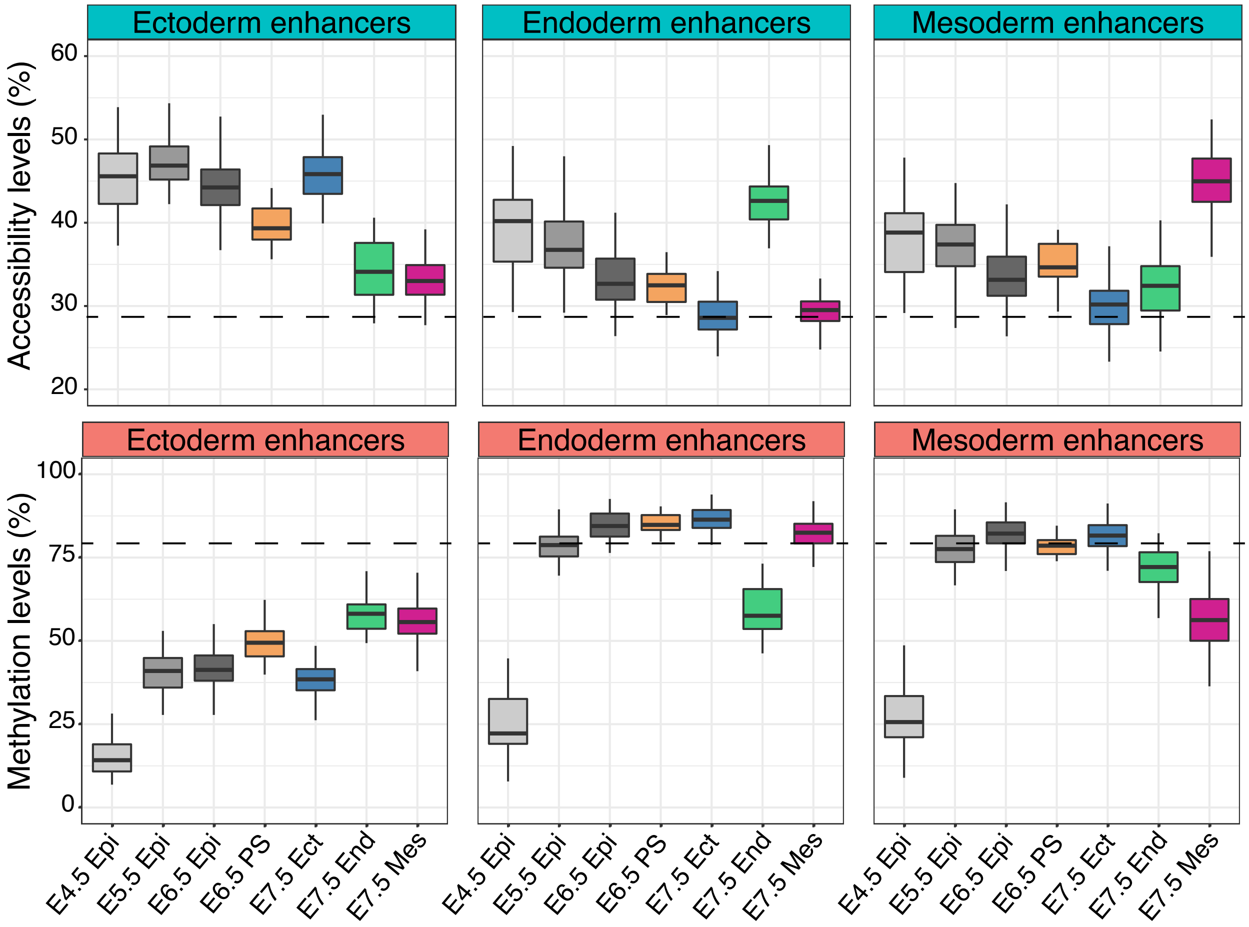


**Figure S24: Epigenome dynamics of lineage-specific enhancers across development.** Boxplots show the distribution of chromatin accessibility (top) and DNA methylation (bottom) of E7.5 lineage-specific enhancers, across lineages and stages. The dashed lines represent the genome-wide averages of GpC accessibility (top) and CpG methylation (below).

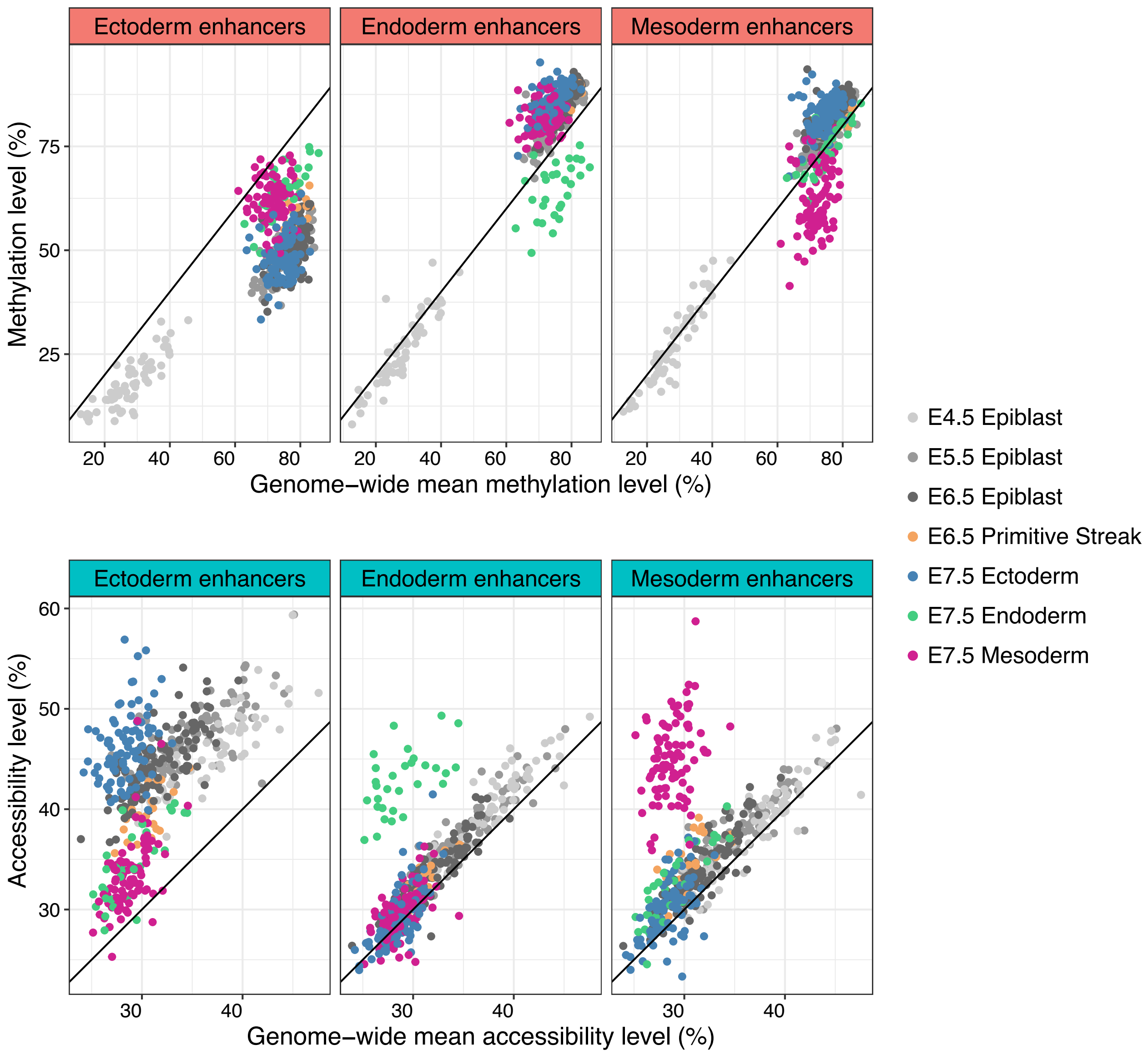


**Figure S25: Ectoderm enhancers are protected from the global DNA methylation and chromatin accessibility dynamics.**

Scatter plots show the genome-wide mean CpG methylation (top) or GpC accessibility (bottom) rate on the x-axis, versus the enhancer-specific mean CpG methylation rate or GpC accessibility rate on the y-axis.

Each dot corresponds to one cell, coloured by the corresponding lineage-developmental stage. The solid line represents the fit of a linear regression that captures the expected genome-wide dynamics. Cells above the line are more methylated (top) or more accessible (bottom) than the genome average. In contrast, cells below the line are less methylated (top) or less accessible (bottom) than the genome average.

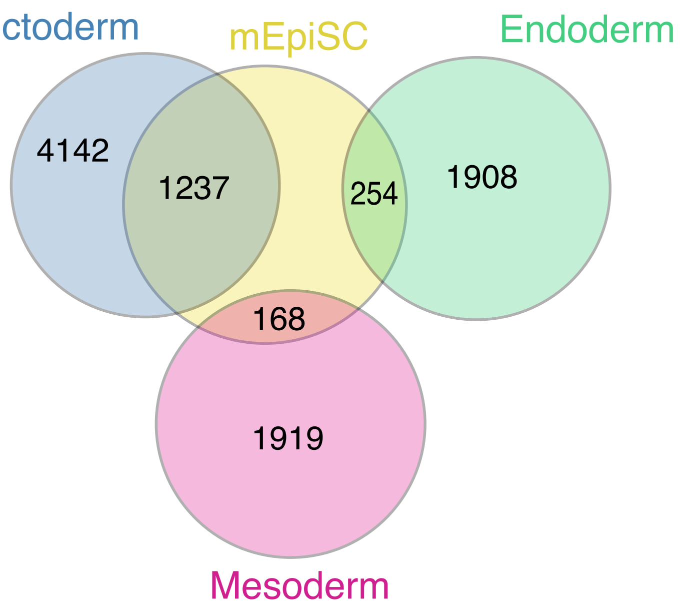


**Figure S26: Ectoderm enhancers overlap with H3K27ac marked regions in mouse Epiblast Stem Cells (mEpiSC).**
Venn diagram showing the overlap between E7.5 lineage-specific distal H3K27ac sites and distal H3K27ac sites derived from *in vitro* mEpiSC[18]. The number of peaks unique to mEpiSC is not shown, as the data was processed with a different computational pipeline, making the absolute number of peaks not comparable.

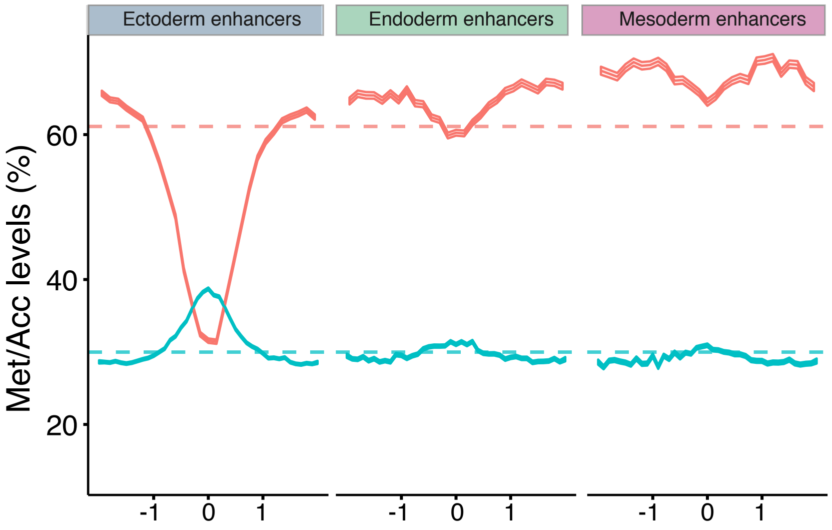


**Figure S27: Ectoderm enhancers are open and unmethylated in mouse Embryonic Stem Cells (ESCs).**
DNA methylation (red) and chromatin accessibility (blue) profiles of E7.5 lineage-specific enhancers in ESCs (processed with scNMT-seq[19]). Shown are running averages in consecutive non-overlapping 50bp windows. Solid line displays the mean across cells and shading displays the corresponding standard deviation.

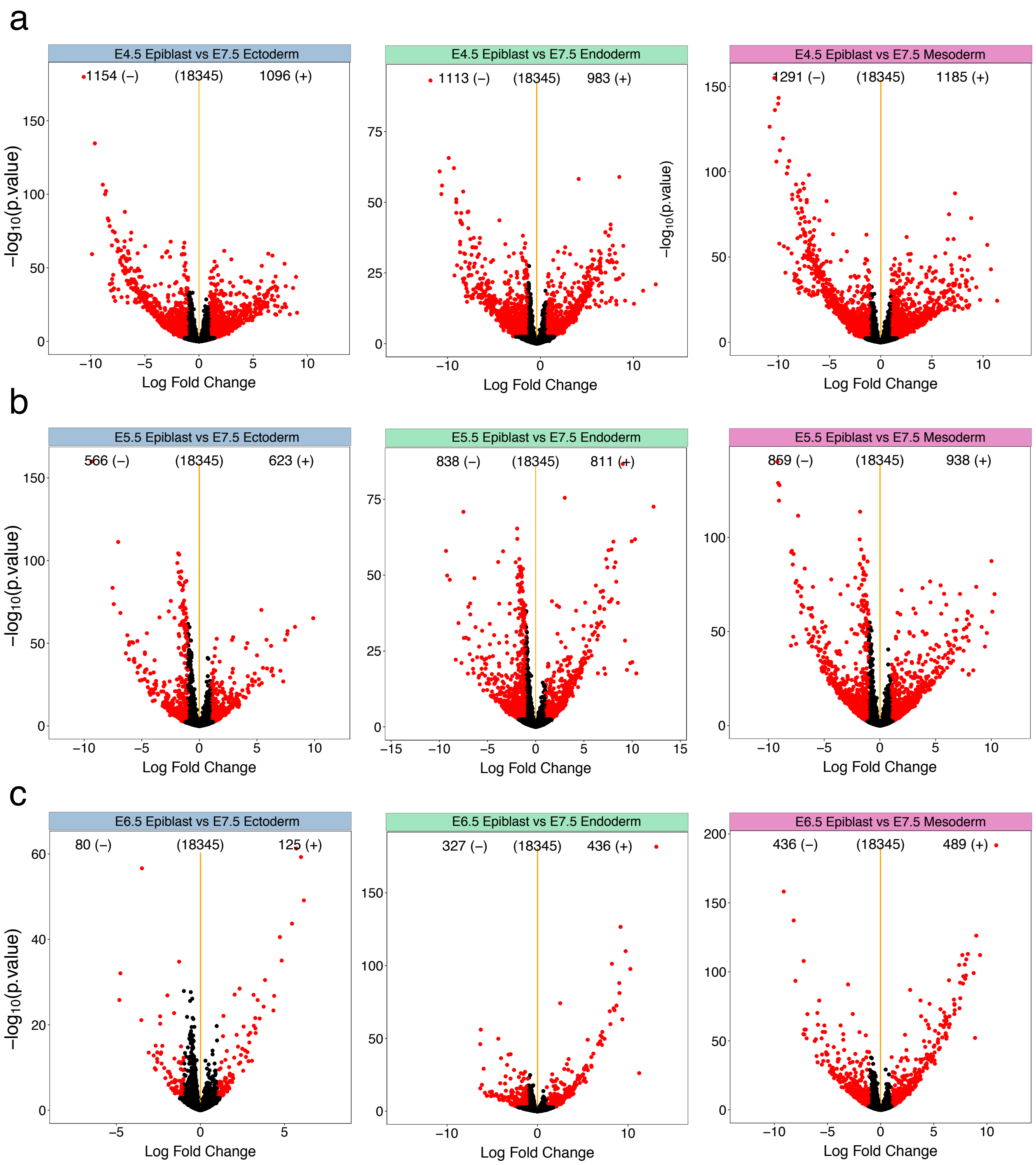


**Figure S28: Epiblast cells are transcriptionally distinct from E7.5 ectoderm cells.** Volcano plots display differential RNA expression levels between epiblast cells from the stages (a) E4.5, (b) E5.5, (c) E6.5, and each of the three E7.5 lineages (ectoderm, left; endoderm, middle; mesoderm; right). X-axis show the difference in mean log2 counts. Negative values for differential RNA expression indicate higher expression in the epiblast, whereas positive values indicate higher expression in the E7.5 lineage. Y-axis shows the Benjamini-Hochberg adjusted p-values. Significant associations (FDR$<$10% and absolute log fold change higher than 2) are coloured in red.

1. Ohnishi, Y. *et al.* Cell-to-cell expression variability followed by signal reinforcement progressively segregates early mouse lineages. *Nat Cell Biol* **16,** 27–37. ISSN: 1476-4679 (Electronic) 1465-7392 (Linking) (2014).
2. Scholer, H. R., Dressler, G. R., Balling, R., Rohdewohld, H. & Gruss, P. Oct-4: a germline-specific transcription factor mapping to the mouse t-complex. *EMBO J* **9,** 2185–95. ISSN: 0261-4189 (Print) 0261-4189 (Linking) (1990).
3. Kalantry, S. *et al.* The amnionless gene, essential for mouse gastrulation, encodes a visceral-endoderm-specific protein with an extracellular cysteine-rich domain. *Nat Genet* **27,** 412–6. ISSN: 1061-4036 (Print) 1061-4036 (Linking) (2001).
4. Mohammed, H. *et al.* Single-Cell Landscape of Transcriptional Heterogeneity and Cell Fate Decisions during Mouse Early Gastrulation. *Cell Rep* **20,** 1215–1228. ISSN: 2211-1247 (Electronic) (2017).
5. Okuda, A. *et al.* UTF1, a novel transcriptional coactivator expressed in pluripotent embryonic stem cells and extra-embryonic cells. *EMBO J* **17,** 2019–32. ISSN: 0261-4189 (Print) 0261-4189 (Linking) (1998).
6. Belo, J. A. *et al.* Cerberus-like is a secreted factor with neutralizing activity expressed in the anterior primitive endoderm of the mouse gastrula. *Mech Dev* **68,** 45–57. ISSN: 0925-4773 (Print) 0925-4773 (Linking) (1997).
7. Saga, Y. *et al.* MesP1: a novel basic helix-loop-helix protein expressed in the nascent mesodermal cells during mouse gastrulation. *Development* **122,** 2769–78. ISSN: 0950-1991 (Print) 0950-1991 (Linking) (1996).
8. Zhang, Y. *et al.* Dynamic epigenomic landscapes during early lineage specification in mouse embryos. *Nat Genet* **50,** 96–105. ISSN: 1546-1718 (Electronic) 1061-4036 (Linking) (2018).
9. Argelaguet, R. *et al.* Multi-Omics Factor Analysis-a framework for unsupervised integration of multi-omics data sets. *Mol Syst Biol* **14,** e8124. ISSN: 1744-4292 (Electronic) 1744-4292 (Linking) (2018).
10. Creyghton, M. P. *et al.* Histone H3K27ac separates active from poised enhancers and predicts developmental state. *Proceedings of the National Academy of Sciences* **107,** 21931–21936. ISSN: 0027-8424 1091-6490 (2010).
11. Liang, G. *et al.* Distinct localization of histone H3 acetylation and H3-K4 methylation to the transcription start sites in the human genome. *Proc Natl Acad Sci U S A* **101,** 7357–62. ISSN: 0027-8424 (Print) 0027-8424 (Linking) (2004).
12. Subramanian, A. *et al.* Gene set enrichment analysis: a knowledge-based approach for interpreting genome-wide expression profiles. *Proc Natl Acad Sci U S A* **102,** 15545–50. ISSN: 0027-8424 (Print) 0027-8424 (Linking) (2005).
13. Ashburner, M. *et al.* Gene ontology: tool for the unification of biology. The Gene Ontology Consortium. *Nat Genet* **25,** 25–9. ISSN: 1061-4036 (Print) 1061-4036 (Linking) (2000).
14. Tamplin, O. J. *et al.* Microarray analysis of Foxa2 mutant mouse embryos reveals novel gene expression and inductive roles for the gastrula organizer and its derivatives. *BMC Genomics* **9,** 511. ISSN: 1471-2164 (Electronic) 1471-2164 (Linking) (2008).
15. Sousa-Nunes, R. *et al.* Characterizing embryonic gene expression patterns in the mouse using nonredundant sequence-based selection. *Genome Res* **13,** 2609–20. ISSN: 1088-9051 (Print) 1088-9051 (Linking) (2003).
16. Bonnafe, E. *et al.* The transcription factor RFX3 directs nodal cilium development and left-right asymmetry specification. *Mol Cell Biol* **24,** 4417–27. ISSN: 0270-7306 (Print) 0270-7306 (Linking) (2004).

17. Finger, J. H. *et al.* The mouse Gene Expression Database (GXD): 2017 update. *Nucleic Acids Res* **45,** D730–D736. ISSN: 1362-4962 (Electronic) 0305-1048 (Linking) (2017).

18. Factor, D. C. *et al.* Epigenomic comparison reveals activation of ”seed” enhancers during transition from naive to primed pluripotency. *Cell Stem Cell* **14,** 854–63. ISSN: 1875-9777 (Electronic) 1875-9777 (Linking) (2014).

19. Clark, S. J. *et al.* scNMT-seq enables joint profiling of chromatin accessibility DNA methylation and transcription in single cells. *Nature Communications* **9.** ISSN: 2041-1723. doi:10.1038/s41467-018-03149-4 (2018).
